# Measuring DNA contents of animal and plant genomes with Gnodes, the long and short of it

**DOI:** 10.1101/2024.10.06.616888

**Authors:** Donald G. Gilbert

## Abstract

Measurement of DNA contents of genomes is valuable for understanding genome biology, including assessments of genome assemblies, but it is not a trivial problem. Measuring contents of DNA shotgun reads is complicated by several factors: biological contents of genomes at species, individual and tissue or cell levels, laboratory methods, sequencing technology and computational processing for measurement and assembly. This compares, and shares, complications with cytometric (Cym) and related molecular measurements of genome size and contents.

There is an obvious discrepancy between cytometric measurements and current long-read genome assemblies (Asm): genome assemblies average 12% below Cym measured sizes, differing in amounts of duplicated content. This report examines five DNA read types to see if they can be used for more precise and reliable discrimination of major genome contents and sizes. The read types are short, accurate Illumina, long Pacific Biosciences, of low and high accuracy, and long Oxford Nanopore Technology of low and high accuracy. Gnodes is the measurement tool used, which maps DNA to assembly, and measures DNA copy numbers for major genome contents of genes, transposons, repeats, and others, using as a measurement unit the single copies of unique conserved genes. Public data of five well studied genomes, human, corn, zebrafish, sorghum and rice, are used for the primary evidence of this work.

Results of this are mixed and open to interpretations: In broad terms, all DNA types measure about the same genome contents, at or below 90% agreement, which is a level that the other complications can contribute. For precision above a 90% level, long read types differ in supporting larger cytometric sizes (low accuracy reads), or smaller assembly sizes (high accuracy reads), with accurate short-reads roughly between. The weight of evidence suggests that low accuracy long reads are less biased for genome measurement, that high accuracy long reads have a bias of reduced duplications introduced by computational averaging or filtering. The several complicating factors noted can produce discrepancies larger than this average Cym - Asm difference, and are a problem to control.

## Introduction

Precision genomics is essential in medicine, environmental health, sustainable agriculture, and biological sciences. Recent efforts with the EvidentialGene project have focused on accurately measuring genome DNA contents with a program called **Gnodes**, a Genome Depth Estimator (Gilbert 2022, 2023). This includes the problematic area of duplicated and highly repetitive structure in genomes, with results that will help inform new and updated genome projects. A genome re-assembly of model plant *Arabidopsis* resolves a 20 year old large discrepancy (Bennett et al 2003). The discrepancy according to Gnodes is missing duplications and repeats, including coding genes, but mostly Satellite DNA, or higher order repeats.

The focus of this paper is to determine how reliable and accurate are DNA read types for measuring genome sizes and their major contents. Genome assemblies are used, but assembly methods are not a focus. Cytometric measures of genome sizes are used for comparisons. For these three basic evidences (DNA reads, assembly [Asm], cytometry [Cym]) it is known that errors of measurement exist and are difficult to identify. Among factors that affect the measurement of DNA samples are biological dimensions: species, population, tissue and cell sources; laboratory and molecular methods; sequencing machine types; computational methods for DNA processing, and secondary methods used for assembly of these DNA samples. Examples of factors affecting measurement are:

a. Biological, within species at levels of population, individual, tissue, and cell. Recent work has solved a genome size discrepancy for Asm/Cym in *Drosophila* species: repetitive heterochromatic regions are under-replicated in thorax and muscle cells, biasing measures and assembly of whole-fly DNA contents (Hjelmen et al. 2020). Maize, the corn plant, has a wide range of genome sizes across populations and inbred lines, from 2 to over 3 gigabases, using Cym and DNA measures (Hufford et al 2021). Most of corn genomes size variation is attributed to active long-terminal repeat transposons (LTR) which comprise 80% and more of these genomes, as discovered by McClintock (1950).
b. Laboratory methods, including biosample processing, DNA extraction. Some examples noted in this work are for "same" plant biosample, one had 40% aphid DNA contamination, the other none; results below detail other lab effects on DNA measures. Methods used to reduce contamination, tissue selections, growth conditions, and others can have effects on genomic DNA contents.
c. Sequencing methods and technology have effects on DNA samples. Early chemistry for short-read Illumina sequencing used PCR-amplifcations that were often very biased; recent PCR-free methods are measurably less biased (Lower et al. 2018). Long-read Pacific Biosciences (Pb) and Oxford Nanopore Technology (ONT) sequencers produce artifacts in homopolymer spans, reported elsewhere and in these results.
d. Computational processing of sequencing data, from base-calling through error-correction, filtering, and other steps in genome assembly and/or measurements. Computational steps that focus on simplifying DNA often come at the expense of removing biologically valid duplicated DNA sequences. Results here show some of the range of these effects, from error correction methods to high fidelity DNA sequences.

These factors can in theory be controlled to improve precision of genome measurements and assemblies. In practice, it is complicated to even learn which factors are important for a given organism, as they interact, and prior knowledge is diffuse, sometimes lost. A prime example is the plant model *Arabidopsis*, where the first assembly at 120 mb was known then to be significantly incomplete (AGI 2000), yet later studies, esp. bioinformatics benchmarks, used this partial assembly as if it were complete (e.g., Vurture et al 2017/GenomeScope, Roach et al 2018/Purge_haplotigs). For this paper, the focus is on c. sequencing methods, with attempts to control a. biological factors, and b. lab methods made by winnowing published data for projects that use the same biosamples and lab methods but vary sequencing methods. Computational post-processing (d) for assembly, e.g. error correction, is not needed here. However the two high accuracy long-read methods reported here, Pb-hifi and ONT-duplex, use computational averaging or filtering, in ways that may bias measures of genome contents.

The lower accuracy forms of Pb and ONT sequencing may or may not provide unbiased measures. Some published low-accuracy ONT and Pb samples are noted as "error-corrected", or filtered against a genome assembly, this may also have introduced a content bias. As this is a complex area of genome research, several DNA measurement approaches have been described (Sun et al. 2017; Vurture et al. 2017; Pucker 2019).

Measurements are based on the basic formula G= L*N/C (Lander & Waterman, 1988) to calculate genome size in bases (G) from observed DNA fragment sizes (L) and number of DNA reads (N), or LN as total DNA bases, divided by a constant C coverage depth of those fragments. The C coverage value is found using unique conserved gene coding sequences as a reliable "inch" for a genome measurement ruler (Gnodes#1). An important assumption of this formula is that DNA fragments are an unbiased sample of nuclear genome. As noted above, this assumption is likely bent or broken for many samples. Sample DNA from non-nuclear sources (contaminants, chloroplasts) need to be measured and subtracted. Measures using K-mer fragment estimation, from peaks of a poisson distribution of fixed-size oligomer sub-fragments of DNA fragments (Li & Waterman 2003) are derived from this same formula, and its assumptions, with added assumptions from a presumed poisson distribution of fragments. Gnodes calculates total genome sizes as well as amount of major contents identified by annotations from genes, transposons, repeats and other sequence features, using coverage depths in DNA samples mapped onto their assemblies, with copy numbers relative to the unit unique gene sequences.

Measures for major contents help to assess the biological validity of results.

## Results

### S1. Short of it

Measurements with Gnodes for both short and long read DNA, with several informatics tools, are reported here. A short result it is that long and short read DNA produce about the same genome size and content measurements, with several caveats about biological, molecular and informatics methods used. This author found no fully reliable set of current methods for genome DNA measurements; any single method may produce very divergent results, versus a median of other methods. However a subset of DNA measurements is more likely to produce reliable results, ones that agree with either cytology measures or assembled DNA measures. A summary of these measures for model and problem species is given in Gnodes#3, Table 1, Measures of genome size where large discrepancy exists.

Genome sizes are measured for five DNA read types, in five species with balanced samples:

- Il or Illu: Illumina short, paired 150-250 bp reads using recent PCR-Free chemistry;
- Oh or Ontd: Oxford Nanopore Tech. duplex (high accuracy), of 2 samples, human and corn [Koren S et al. (2024)]. Computational averaging and/or filtering is used to achieve accuracy;
- Ol or Ontl: Oxford Nanopore Tech. low accuracy, long reads;
- Ph or Pbhi: Pacific Biosciences high-fidelity or CCS reads. Computational averaging and/or filtering is used to achieve accuracy;
- Pl or Pblo: Pacific Biosciences low-fidelity or CLR reads.

Genome assemblies published from these DNA read sets are used for measuring read coverage and copy numbers, with annotations of genome contents for transposons, coding genes, rRNA genes, satellite repeats, tandem repeats and other as produced by those projects or this project. Non-nuclear DNA has been removed (chloroplast, mitochondria and contaminants identified by NCBI SRA reports).

#### DNA measures of genome sizes

Figure F1.sum.A summarizes results with respect to five DNA read types (Illumina short, Oxford Nanopore and Pacific Biosciences, low & high accuracy of these two long types), and assemblies, of five well studied animals and plants. This figure plots nuclear genome sizes as measured with Gnodes [Gnodes#1] relative to cytometric measured sizes. Species genomes examined here in detail are human (3200 megabases), corn or maize (2600 mb), zebrafish (1600 mb), sorghum (800 mb) and rice (450 mb), for which recent assemblies using long-read DNA are available (2020-2023), and for which a balanced set of read types for the same assembly bio-sample, generally from the same project, are publicly available.

**Figure F1. sum.**
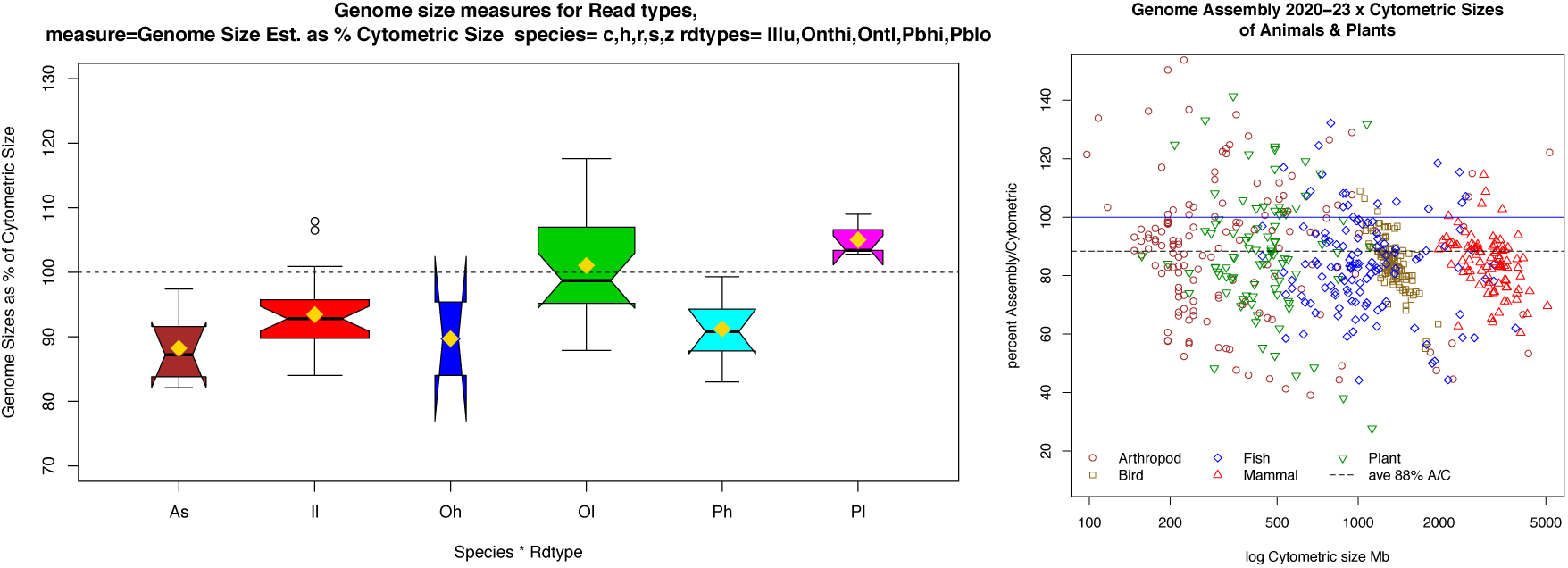
**(A)**. Genome size estimates relative to cytometric size (Y) for assemblies (As) and five read types (Il, Oh, Ol, Ph, Pl), average of five species (human, zebrafish, corn, sorghum and rice). These box plots show median (black center line), average (gold diamond), and ranges. Box width is a sample size clue, where Oh with 2 samples is narrow. The horizontal dotted line at 100% is cytometric measure. The average close to median suggests normally distributed values. Details per species are below. **(B)**. Genome sizes for 500 animal and plant species, as Assembly / Cytometric size percentage (Y) in relation to cytometric sizes (X-axis in log megabases). Colored symbols indicate the clade of arthropods, birds, fishes, mammals and plants. Horizontal lines are cytometic average (100%) and assembly average (88%) (Figure I1 of Gnodes#3 doc).

**Figure F1. sum.C.**
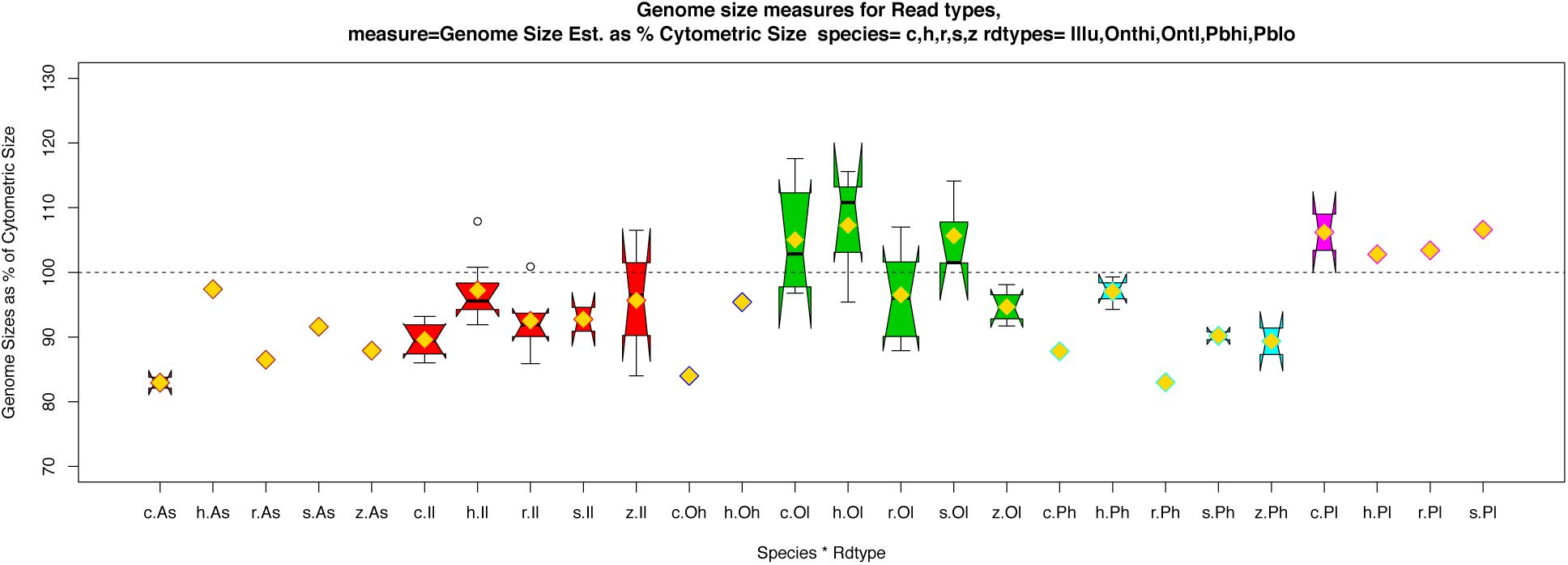
Genome sizes measured for DNA types (Il, Oh, Ol, Ph, Pl) and assemblies (As) of five species (c=corn, h=human, r=rice, s=sorghum, z=zebrafish). Averages are marked with gold diamond in these boxplots, sample size indicated by width of box (diamond only for 1 sample).

**Figure F1. sum.D.**
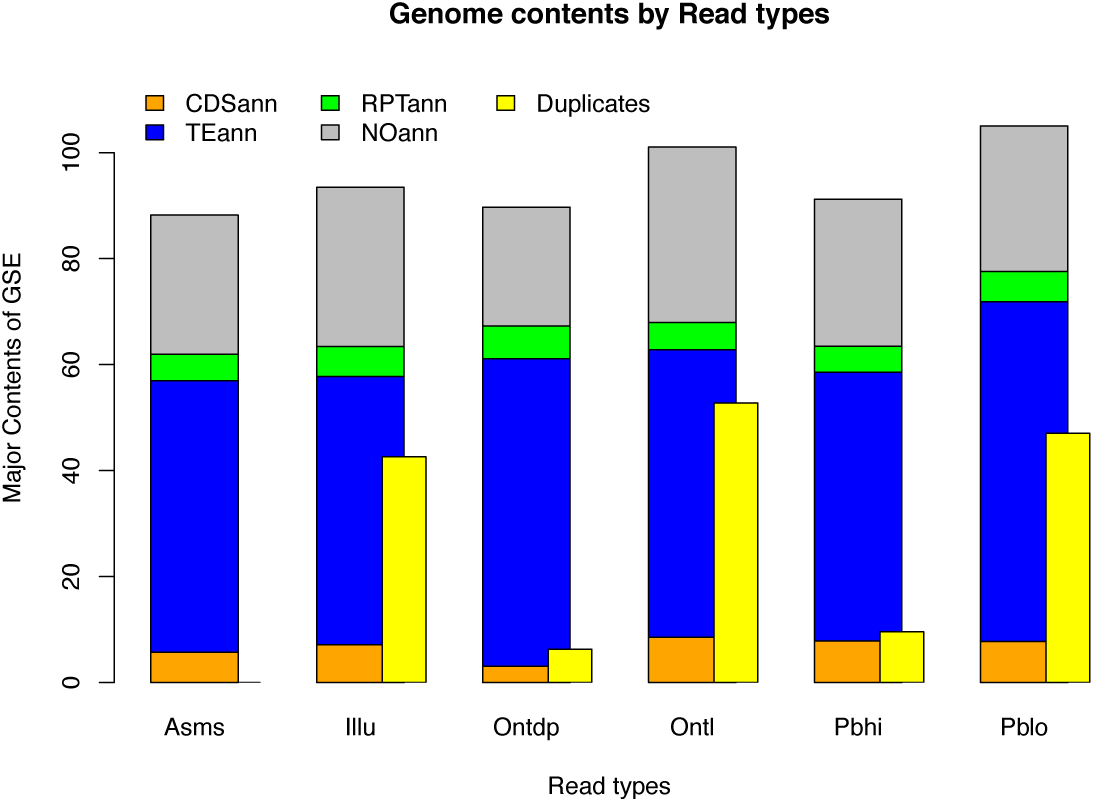
Major contents to genome sizes measured for DNA types and assemblies, averaged over species. Contents are shown here as percentage of total size relative to cytometric sizes (averages in F1.sum.A figure), so that Pblo and Ontl are at/above 100% total, while Asm, Ontdp and Pbhi are at/ below 90% total. Contents annotated are gene CDS (orange), transposons (TE, blue), and repeats (RPT, green) including rRNA, simple and high order repeats, with NOann the remainder (gray). Duplicates over all contents (yellow) are the multi-mapping read percentage for DNA types.

There is often a difference in genome sizes measured from cytometry and assembly, assemblies are on average smaller. Figure F1.sum.B illustrates this, comparing recent (years 2020-2023) genome assemblies of animals and plants, versus their cytometric measured sizes, for about 100 species each of arthropods, birds, fishes, mammals and plants. A majority of assemblies are from 15% to over 50% below cytometric sizes. Assembly sizes average 88% of cytometric sizes for the five species in A, the same as for the assemblies of 500 species of B.

These summaries indicate the assemblies are below cytometric measures, though human is close to 100%, the rest range to and below 80%). Three DNA types measure below Cym, nearest to assembly, one is well above Cym and others, and one, Ontl is rather close to Cym sizes, on average. The average shown here is dependent on species sampled, and the many other factors (lab and info methods). Both high-fidelity types (Ph and Oh) average at 88%, Illu is 92%, and Pl, Pblo is notably above 100%. Only two ranges cross Cym midline, Ontl and Illu, though ranges are roughly proportional to sample size.

Thus the question, which is most accurate, or can that even be determined? DNA from the same bio- samples and projects measure differently enough that they clearly disagree which of assembly or cytometric sizes are the more accurate.

Figures F2.rRNA (A,B) give examples of these discrepancies, for the highly duplicated ribosomal RNA genes, where several DNA types and genome assemblies are measured for gene copy numbers. These figures show copy numbers as measured across the spans of large subunit rRNA gene (3000 to 5000 bp). A human genome assembly has about 230 copies, though some types of DNA sequences measure at 300 copies. In contrast, a corn genome assembly has 400 copies of rRNA, but most DNA types contain an order of magnitude higher at 3,000-4,000 copies. This and related measures of DNA suggest that corn and many other recent genome assemblies are still missing large numbers of duplicated genes and other contents.

Agreements for high copy genome contents are examined, for the five DNA read types, in three species with balanced samples of these read types. Agreement here means two or more read types have similar high copy estimates for genome spans. Agreement is an uncertain proxy for truth, where no certain measure of high copy genome contents is known. Agreement summaries are plotted in Figures F3.agr (A,B) with a 3-number glyph showing percentages of high-copy amounts (HC) relative to assembly, those with agreements (AG) relative to high-copy counts, and agreements relative to any-agreement subset (AA). In example figure F2.rRNA, copy agreement of Pblo and Ontl exists for both species, while Pbhi and Ontd disagree for one, which lowers AG and AA values. Figure F3.agr.B panel for rRNA shows these agree values.

**Figure F2. rRNA.**
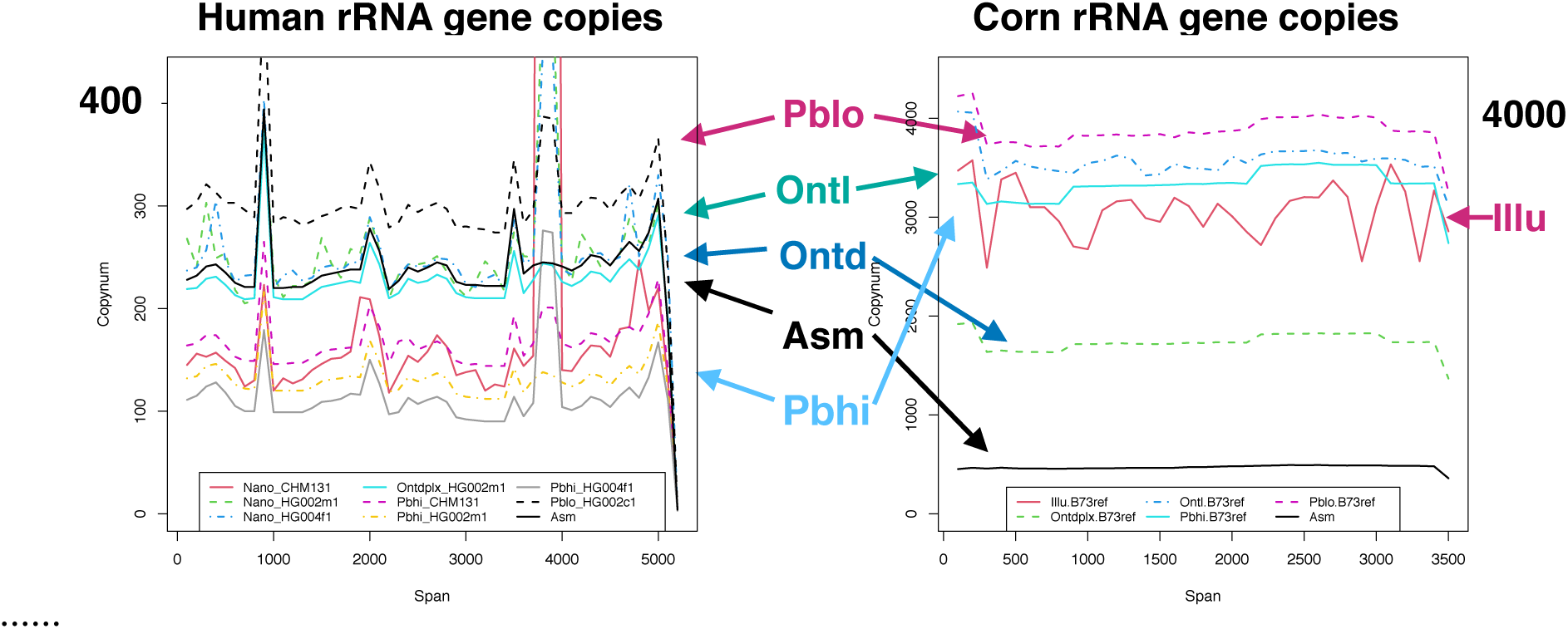
Large subunit rRNA gene copies as measured in DNA read types and assembly, over gene spans of 3000-5000 bp, for (A) human with maxima around 400, and (B) corn, with maxima of 4000 copies.

**Figure F2. rRNA.**
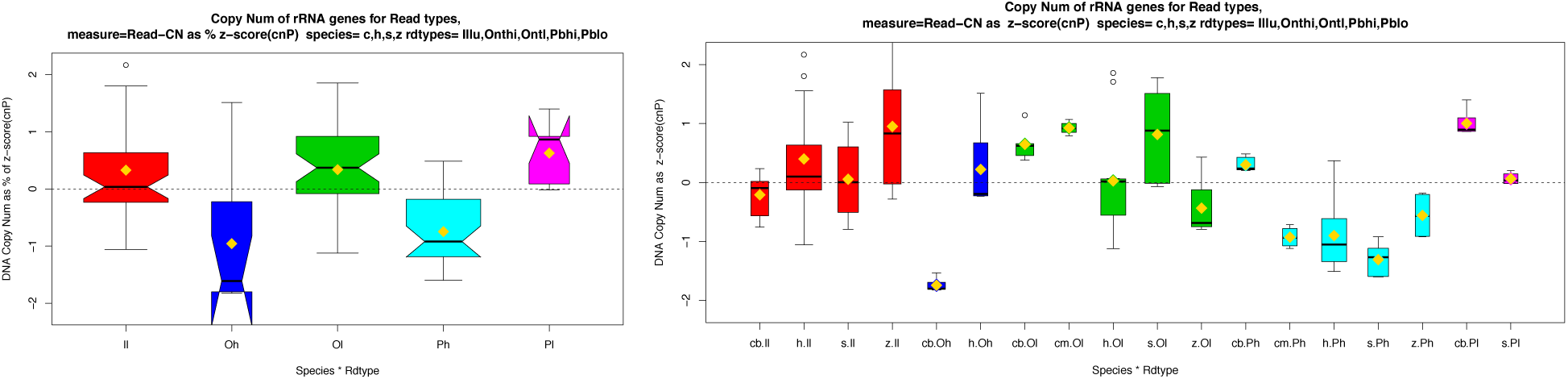
rRNA copy number as species-relative z-scores (y-axis), by read type averaged over species (C) or for each species (D).

**Figure F3. agr.**
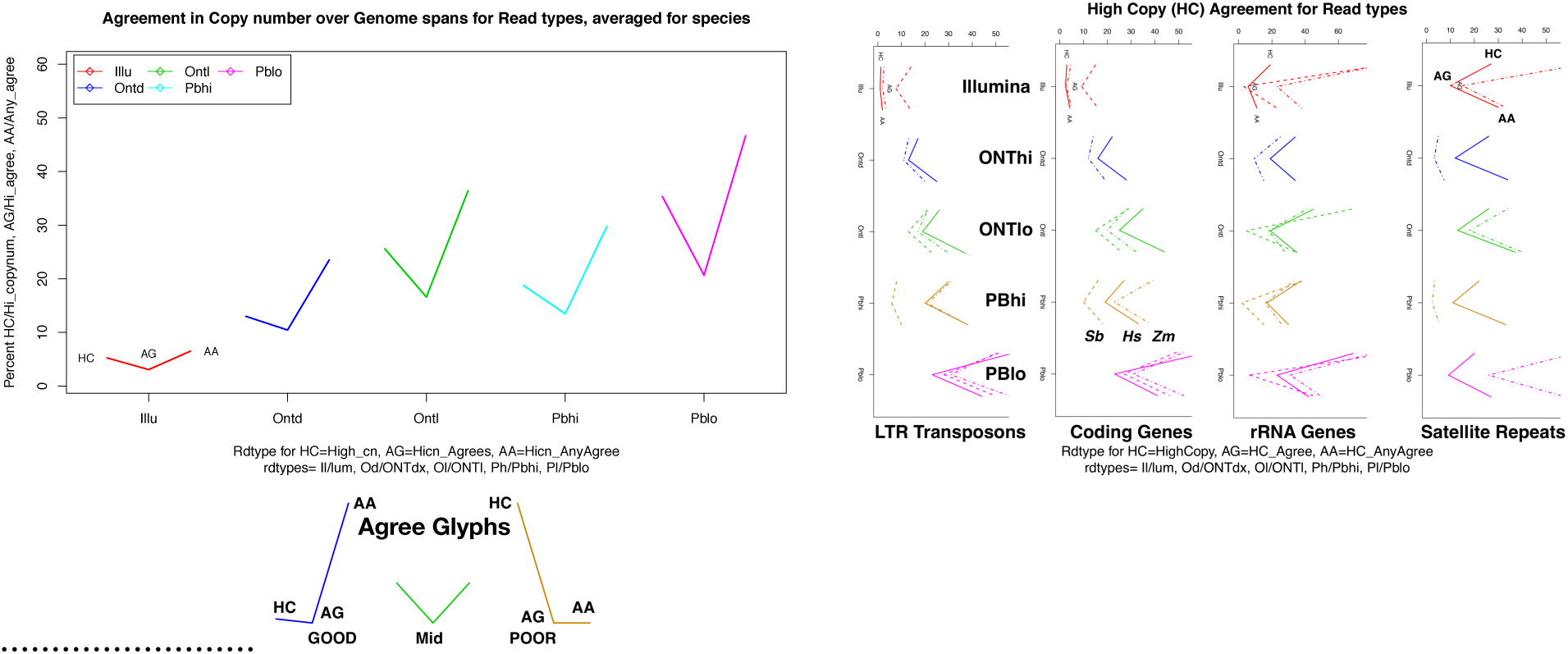
Agreement in high copy number measures among read types, for (A) average over species and contents, and (B) by major contents of LTR transposons, coding genes, rRNA genes and satellite repeats. Agree glyphs and details are described in section S2e.

Over these high copy genome content types, Ontl has the most consistent agreement. Pblo has evidence of excess in false high copy, along with more numerous HC spans. Pbhi, and the small sample of Ontd-duplex, show more variability over species; the human samples are similar to Ontl, but the plant samples have some very low agreement in high-copy spans. HC is lowest in Pbhi and Ontd samples, i.e., they match copy numbers in the genome assemblies. Accuracy of duplicated spans is uncertain for these genome assemblies, but evidence cited point to under-assemblies in such spans. Illumina short reads have lowest HC over all, and low agreement.

Although mapping DNA reads to Asm is affected by algorithms and other factors, this is not an important part of DNA measures with Gnodes. The important map measure of unique conserved genes coverage is accurate, due to uniqueness. It is however necessary to properly categorize DNA bases, and for that one must classify un-mapped and poorly mapped portions. Measuring contamination (non-nuclear DNA) is essential, as the remaining unmapped bases are genomic contents, or artifacts. Long read DNA introduces a question of unplaced bases in long reads that are partly mapped to assemblies.

Though minimal in short read data, these can account for 5% to over 20% of some species long read data.

These unplaced DNA spans, when extracted and re-aligned to their assembly, will mostly re-align at duplicated regions (Table T1). Although sequencer artifacts can produce spurious inserted bases (homopolymer effect known especially for Pblo), those have been measured as short insert spans of 2-10 bases, adding 1%-2% to sampled DNA.

**Table T1.**
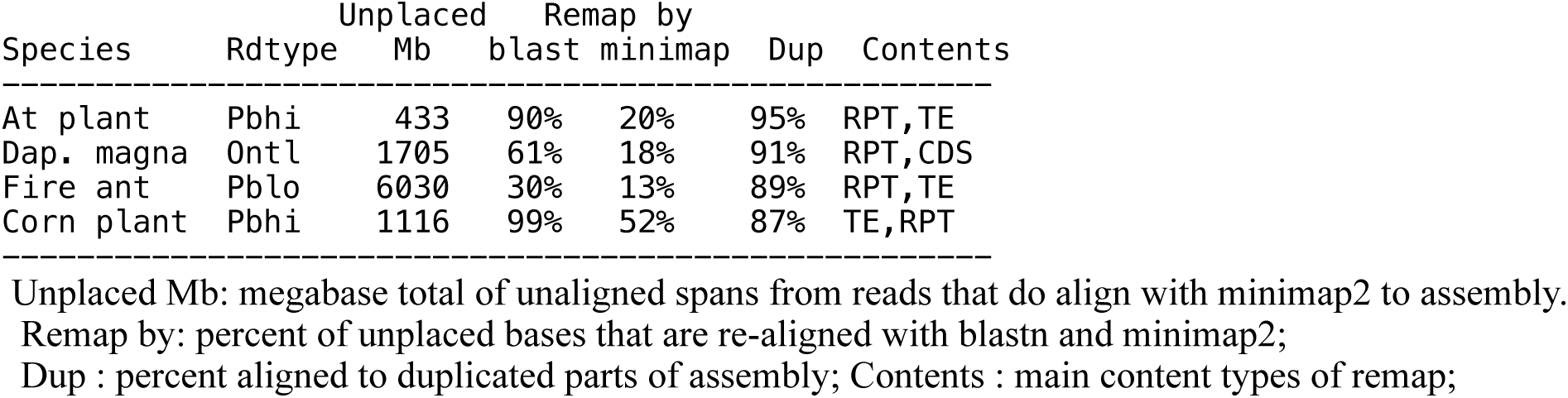
Re-map rates and contents of long-read unplaced spans (details in Table T3)

### S2. Long of it

When this project first examined DNA types in 2022, a result was that read types produced roughly the same genome sizes and content estimates. That work spanned small genomes (150-250 Mb) of arabidopsis, fruitfly, daphnia and others to middle sized genomes (1000-3000 Mb) of human, corn, chicken. Useful were two projects of arabidopsis ecotype lines that included Cym measures, long and short DNA samples, offering a gradation of comparable data of one species. As summarized in Table T2.A, the difference measured for long - short reads is insignificant.

**Table T2.**
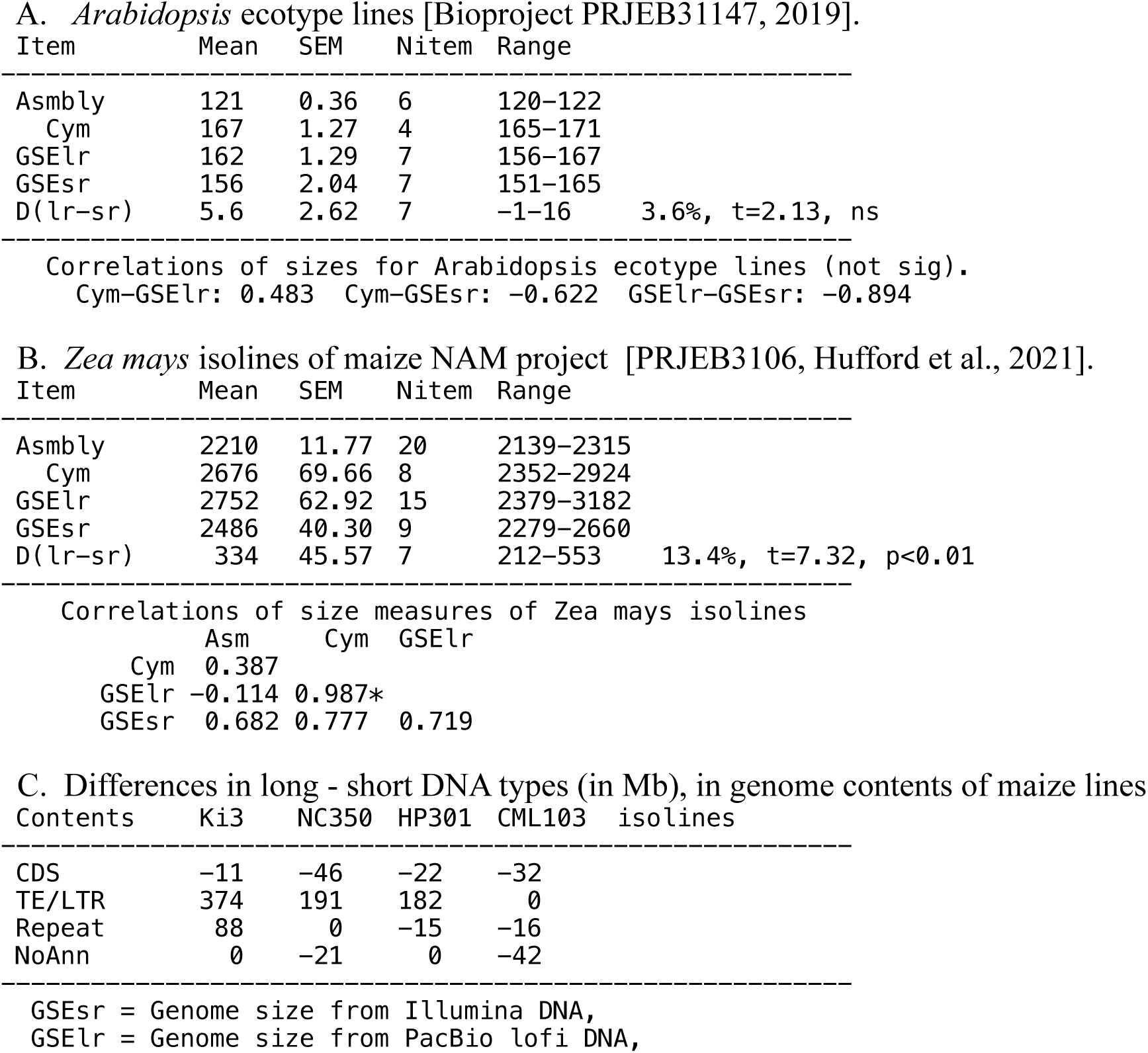
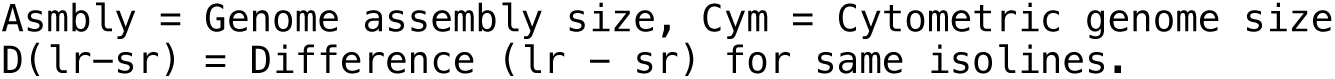
Genome sizes (Mb) for isolines of two plants, from assembly (Asm), flow cytometry (Cym), short (GSEsr) & long (GSElr) DNA.

Some discrepancies to this simple answer were seen for larger genomes. That led to further examination of results from the Maize NAM isolines, where Cym measured a larger range in genome sizes (2,000 Mb to 3,000 Mb). One study (Hufford et al., 2021) of corn isolines provides short and long reads, and their assemblies, for a balanced, same-project data set. Gnodes measured a significant long-short read discrepancy: 334 Mb or 13% of genome sizes (Table T2.B).

Examination of genome contents where lr - sr differ in corn isolines suggests that transposons, which comprise 80% or more of these genomes, contain most of this discrepancy, along with repeats (Table T2.C). The isolines sampled had large variability in Cym sizes, TE content, and the DNA type difference. Further investigation was needed to see if this long-short discrepancy is a consistent read property, or some other methodological artifact.

#### S2b. DNA measures of genome sizes by species

The average shown in F1.sum.A is dependent on species sampled, and the other factors (lab and info methods, see Introduction). Figure F1.sum.C shows boxplots of the five species separately, where means and medians across species are clustered about each DNA type’s average, suggesting the DNA type average is a valid result.

Major contents can differ markedly across species, although this depends on type and quality of these annotations. Transposon portions range from 35% in rice, to 50% in human, 60% in sorghum and zebrafish, to over 85% in corn. Sorghum (800 Mb) and zebrafish (1600 Mb) genomes have similar proportions of CDS (9%) and TE (60%). Examining these contents for effects of read types is difficult even for one species. There are no large differences over read types when averaging over species, as figure F1.sum.D shows. For transposons, Pblo has a slight but not significant larger portion than for other types. CDS is roughly constant percentage, though it is very species/size dependent, as the smallest genome here, rice has a higher portion as CDS balanced by its much lower TE portion. The Ontdp values are skewed due to a sample of only corn and human, but is roughly consistent with the other high fidelity type, Pbhi.

Duplicates differ greatly by DNA type, seen in figure F1.sum.D, but this is difficult to interpret, as alignment algorithms affect this: short reads (Illu), and low fidelity long reads (Ontl, Pblo) multi-map more often due to read structure. However, the higher duplication percentage for Ontl, Pblo, and Illu coincide with their larger genome size estimates, as the size measures differ due to duplications rather than unique DNA contents.

#### S2c. DNA measures of rRNA copy numbers

Copy number of rRNA genes varies greatly, between and even within species (Figure F2.rRNA above for examples of human and corn), and is often under-assembled due to many high-identity duplications. As well, DNA read measures of rRNA abundance are known to be error-prone (Morton et al. 2019). The results for rRNA copies are given here as z-scores calculated for each species, to show read-type effects among species, rather than relative to Cym or Asm counts (Figure F2.rRNA below). The two high-fidelity long-read forms, Pbhi and Ontd, have lowest copy counts of rRNA, Pblo is the highest count, and Ontl and Illu have middle values. This trend holds across species, though the two samples for Ontd are too limited for a conclusion.

#### S2d. DNA measures of Transposon copy numbers

Copy number for transposons does not vary greatly across species, nor over DNA read types, on average. This lack of variation might be due differentiation of transposons, or low sequence identity. The box plots (Figures F4.TEcn) show a large range of transposon values for each sample, i.e. some have high identities. For corn, with the most LTR transposons, the problem noted above (Table T2) shows in box plots where corn Illu has fewest copies, while corn Pblo has the most, among the species and DNA types sampled here.

To see if DNA copy numbers differ for transposons with high identity, all high-identity chromosome segments (hiids, spans of >2000bp with >98% identity) were marked, and DNA copyn of transposons on those segments measured. For corn and sorghum, this subset showed almost no change in average copyn of the DNA read types, from totals of Figure F4.TEcn. The largest change, of 1-2%, was for increased Illu copyn on hiids segments, possibly a map-method effect. High identity subsets comprised about 10% of all transposon spans.

**Figure F4. TEcn.**
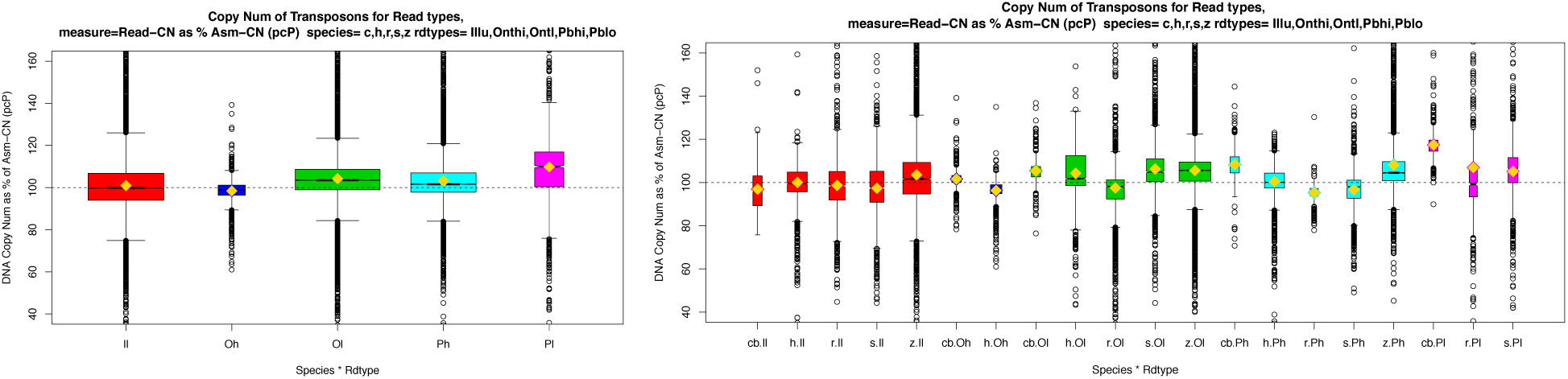
(A,B). Transposon copy numbers as box plots, by read types (A) averaged over species, and showing each species (B). Copy numbers here are relative to assembly copyn (y-axis, 100% matches assembly). Box plot glyphs and x-axis labels of read types and species are as for F1.sum figures.

Example LTR transposons that show copy variation among read types, for corn and sorghum, are pictured in Figures F5.TEx (A,B,C,D,E,F). One result from examining individual LTR cases is that read types are inconsistently different in measuring LTR copy numbers.

**Figure F5. TEx.**
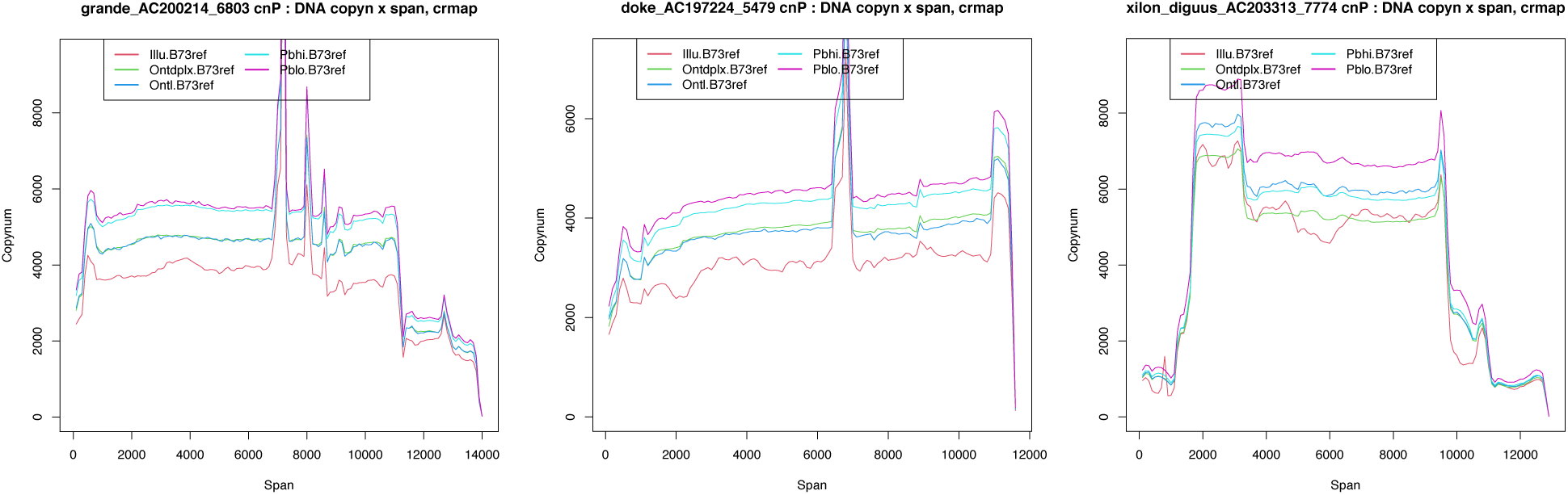
(A,B,C). Corn transposon spans showing copy number (Y) over span (X) for 5 read types of 3 TE examples.

**Figure F5. TEx.**
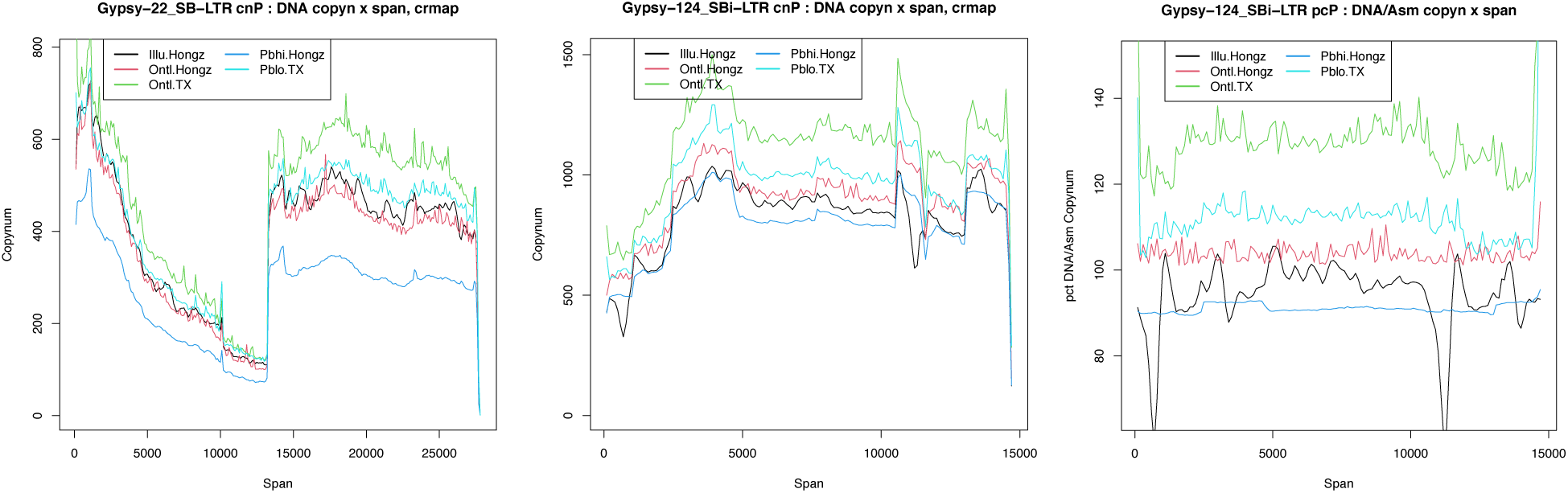
(D,E,F). Sorghum transposon spans showing copy number (Y) over span (X) for 5 read types of 3 TE examples.

**Figures F6. agrd.**
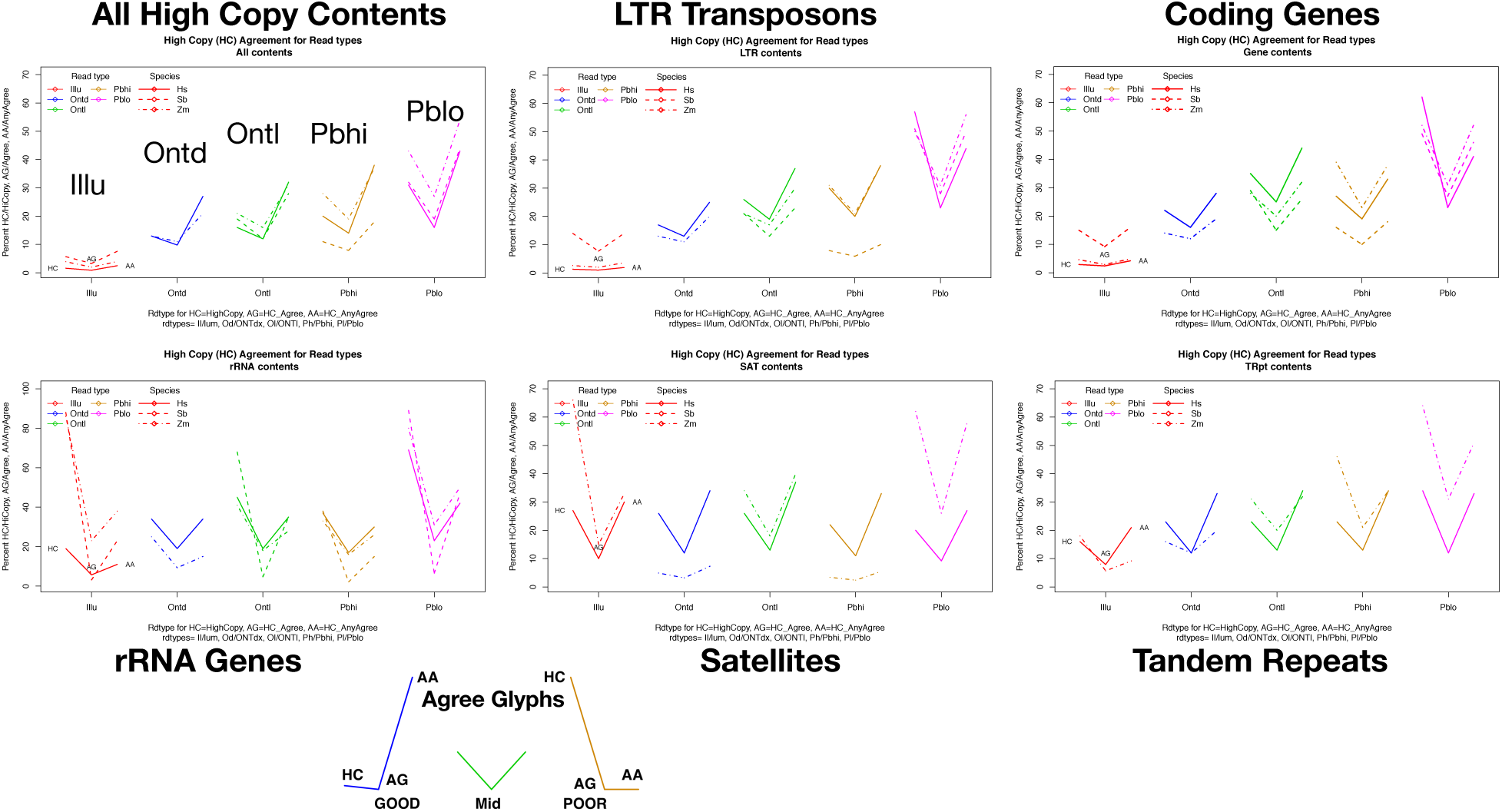
(A,B,C,D,E,F). High copy agreement measures for read type over genome content types. Only contents with copy numbers above 1.5x are measured, e.g. duplicated coding genes and transposons. All high copy contents (A) includes or averages all of these contents. Satellites (E, from RepeatMasker results) and tandem repeats (F, from tandem-repeat finder results) have similar, overlapping content but are annotated differently, and have somewhat different measures.

#### S2e. Agreement for high copy genome contents

Agreements for high copy genome contents are examined, for the five DNA read types, in three species with balanced samples of these read types. Agreement here means two or more read types have similar high copy estimates for genome spans. Agreement is an uncertain proxy for truth, where no certain measure of high copy genome contents is known.

The base-line of one copy derived from unique conserved genes is founded on evolutionary observation, with some molecular understanding, that certain genes are never duplicated. The other end, or top-line, of multiple copies is properly represented by ribosomal RNA genes that are highly duplicated in animals and plants (Hall, Morton & Queitsch 2002). However counting rRNA genes to get a top-line is imprecise, partly due to the flucuations observed in rRNA copies, within species, and likely within individuals. Other genome contents that have many copies are also known to flucuate, so that measures on one biological sample tend to disagree with other samples of the same species, iso-line, individual. Given the one copy base-line for DNA samples from unique genes, higher copy numbers can be estimated, with uncertainty that depends on whether there is a linear relation from one to N copies, and whether there is uniform coverage depth in DNA read samples.

Agree Glyphs draws examples of three percentages of agreement, HC, AG, AA, for high-copy genome spans, with "good", "poor" and middle examples.

- HC counts high-copy spans for this read type (HCi), relative to HC spans for any read type (HCa), where HCa is same denominator for all read types, or HCi/HCa.
- AG is Agreement for high copy spans, counts read spans that are in agreement with another read type for high copy (AGi) relative to all HCi counts, or AGi/HCi. AG is analogous to specificity, i.e. true positives / (true pos. + false pos.)
- AA is AnyAgree, counts of read spans that agree with spans where there is agreement of samples, or AGany (AGi numerator counts agreement for this read type, and AGa denominator counts spans with 2+ agreeing read types). AA is analogous to sensitivity, i.e. true positives / (true pos + false negatives).

A good agreement glyph (backward L) has large AG, relative to HC, and large AA; a poor agreement glyph (forward L) has small AG relative to HC, and small AA. The observed cases are mostly in between. rRNA genes, and Satellites, with extreme high copy, has some poor L cases for Illu and Pblo. LTR transposons and coding genes have good-ish numbers especially for Ontl read type.

Over these several high copy genome content types, Ontl has the most consistent agreement. Pblo has evidence of excess high copy, along with its generally more numerous HC spans. Pbhi, and the small sample of Ontd-duplex, show more variability over species; the human samples are similar to Ontl, but the plant samples have some very low agreement for high-copy spans. HC is lowest in Pbhi and Ontd high-fidelity read samples, that is they match copy numbers in the genome assemblies. There is no certainty on accuracy of the duplicated spans in these genome assemblies, but the evidence cited here point to under-assemblies in such spans. Illumina short reads fair poorly, with lowest HC over all, and low agreement. It is not clear to the author if this is due entirely to the short read data, or may be partly an artifact of how read copy numbers are measured on genome spans, though effort was made to test in various ways and measure in an unbiased way.

#### S2f. Mapping Long & Short DNA to Assemblies

Several long-read DNA mapping programs were tested (see Methods): minimap2, Winnowmap2, blend, lra (long-read-aligner), TandemTools. BWA and minimap2 were used for short-read DNA mapping. minimap2, widely used, is the default choice for Gnodes, with others as options. In general, error-full long reads (pacbio-lofi and oxford nanopore) are less accurately aligned and measured for coverage depth compared with highly accurate short-read DNA of current Illumina sequencers. This is a result of hundreds of Gnodes analyses across the range of animal and plant genomes. In this project, winnowmap produced small differences from minimap2 which were inconsistent in measurement of repetitive content. Winnowmap2 was designed to address the problem of inaccurate repeat mapping, and it is possibly a more accurate approach, with mistakes in this author’s tests of it (Table T6_read_mappers_long_short).

For mapping errors in figure F7.MErr, results are averaged for 6 species, done using minimap2 mapping. Results are as expected from prior work. Total DNA bases in read sets, minus any contaminants, is the percent denominator. Numerator base counts of these are: unAlign counts total - aligned (including mismaps), unPlaced counts unaligned bases from reads that are partly mapped, unMap counts bases from unmapped reads, Insert counts read bases marked as insertions, Delete counts assembly bases with no read bases, and Mismap counts mismatches in alignment. Unaligned is the sum of unplaced and unmapped DNA. The unplaced value is often overlooked, but is a significant amount in error-ful read types Ontl and Pblo. The sample size per species is too small to detect species x read-type effects. A few possible effects are likely laboratory method artifacts.

**Figure F7. MErr.**
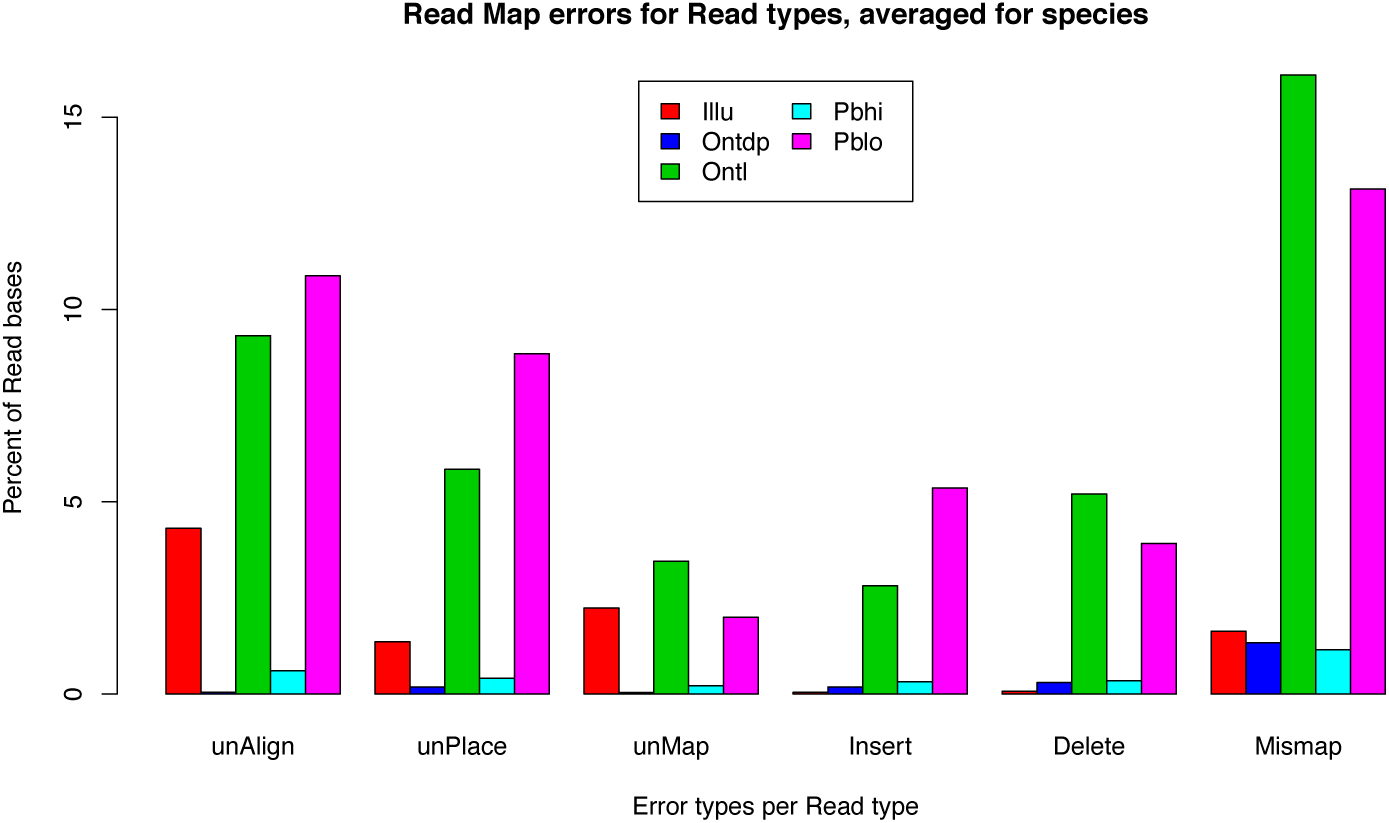
Mapping errors by read types Illu, Ontd, Ontl, Pbhi and Pblo. Error types are described in text; unAlign, unPlace and unMap are measures of total genome content.

While mapping methods affect alignment measures, this is not too important for genome size and contents measures with Gnodes. The essential measures are unique genes coverage, for the baseline Cucg value, total read bases with the assumption these have unbiased coverage of the genome, and measures of any contaminant DNA contents to be subtracted. The unplaced and unmapped bases add to size and content measures. For chromosome assemblies of reasonably good quality, only duplicated spans have problematic alignments.

#### S2g. Measuring high-copy duplications is difficult and inaccurate

Long-read DNA mapping introduces a problem of interpretation where parts of the long read do not map to an assembly, and other parts do. This is common in repetitive spans, where all bases of a long-read can be interpreted as belonging to a genome. These "unplaced" read bases are part of Gnodes analyses, which are placed for practical measures at the point of last mapping. With short-reads, unplaced bases are a minimal amount (1%). For long-reads these are significantly higher: 5% to 15%, and can be over 20% for some long-read data (fire ant HOR spans for example). These unplaced DNA spans, when extracted and re-aligned to their assembly, will mostly re-align at duplicated regions (Table T3).

**Table T3.**
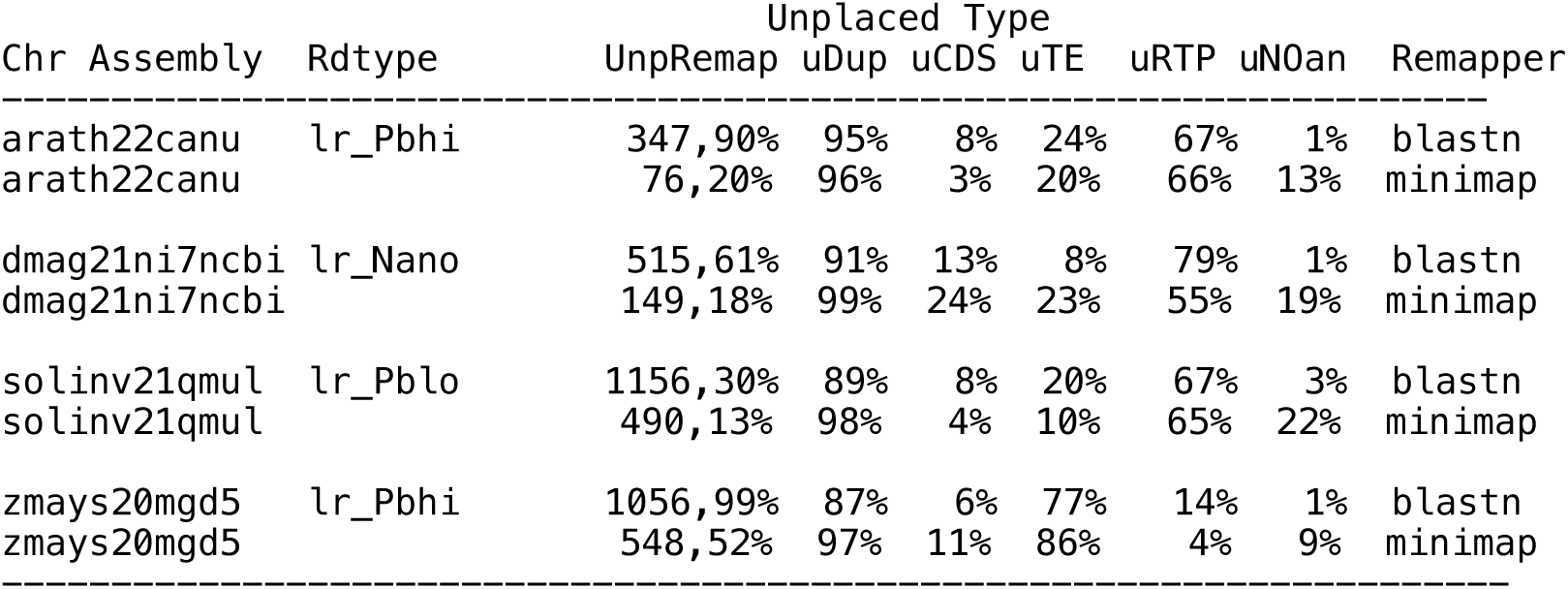
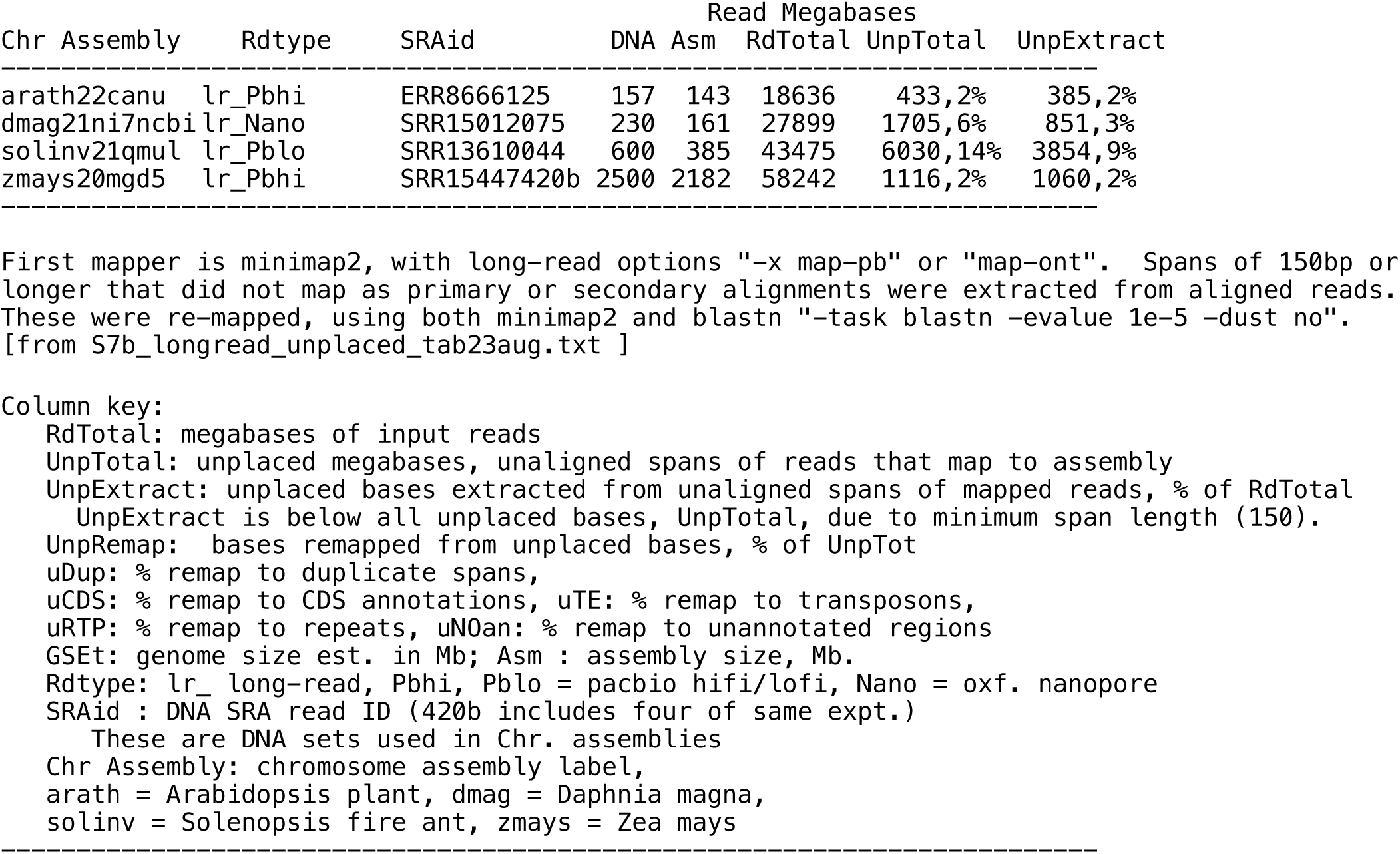
longread_unplacedbases. Unplaced sequence spans from long reads that partly map to chromosomes were extracted and re-mapped. Megabases aligned to annotated content types are indicated, see column key. Unplaced spans of long reads contain duplicated content, the main type depends on species, e.g. telo- and centro-mere repeats in Arabidopsis, Daphnia and fire ant, and transposons in Zea mays. Gene coding sequences are also unplaced in these species.

MGSE (Pucker 2019), another map-based estimator, undercounts DNA genome sizes and has larger error than Gnodes, due to methods that reduce duplicate DNA measurement. These reductions are informative for understanding DNA sample measures: a. only bases mapped to assembly are counted by MGSE, ignoring significant numbers of unplaced bases on reads that map partially, a bias against mis- and under-assembled regions; b. reads that do not map to chromosomes, but are not measured as contaminants, are not counted; c. biological duplicate reads were filtered out by the author, with Picard MarkDuplicates that removes all but one of co-mapped reads, a bias against full depth measure of under- assembled spans; and d. simpler, less precise method of measuring Cucg, depth of DNA at unique genes.

The amounts of genome DNA in these partitions (a, b) are significant, and need to be accounted for when measuring accuracy of genome assemblies, as they putatively identify inaccuracies. Table T4 summarizes these, along with measured contaminants for one study, the *Arabidopsis* ecotype samples of Van De Weyer et al 2019. Measure of aligned bases are nearly the same for Gnodes and MGSE. MGSE as calculated here used the same gene loci for CU depth measure (d), and was not reduced by removing duplicate reads (c). Unplaced bases from mapped reads add a significant 10% to the sizes, and are not accounted for by sequencing adaptors or artifacts, so can be classified as genomic DNA. Unplaced bases are greater in duplicated regions. Unmapped reads account for 4% of the better DNA samples, after removing measurable contaminants (bacteria, adaptors, endogenous non-nuclear DNA). Although this class is ambiguous, there is no positive evidence to exclude it from organismal genomic DNA; this author recommends regarding it as such until evidence warrants otherwise.

**Table T4.**
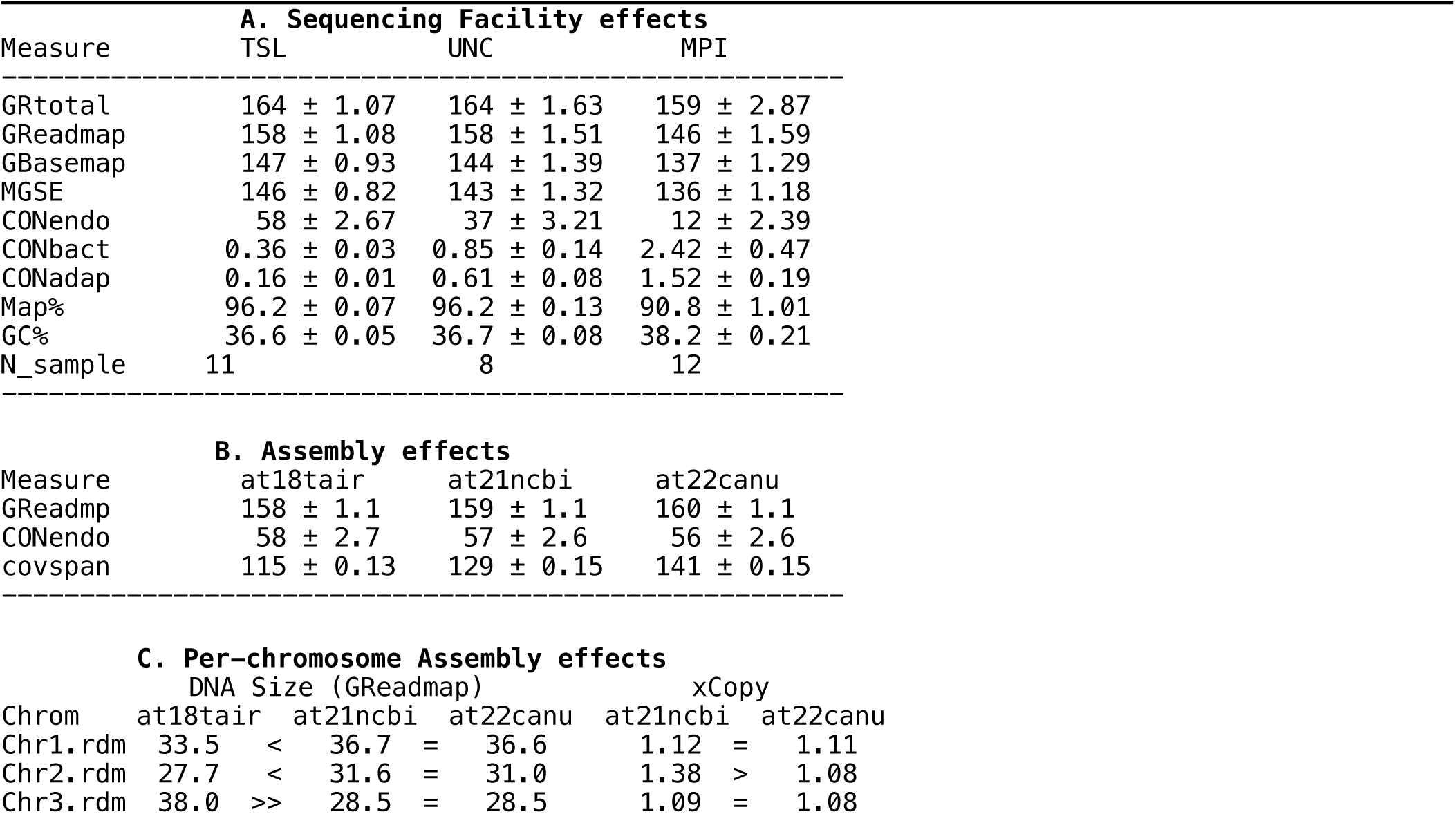
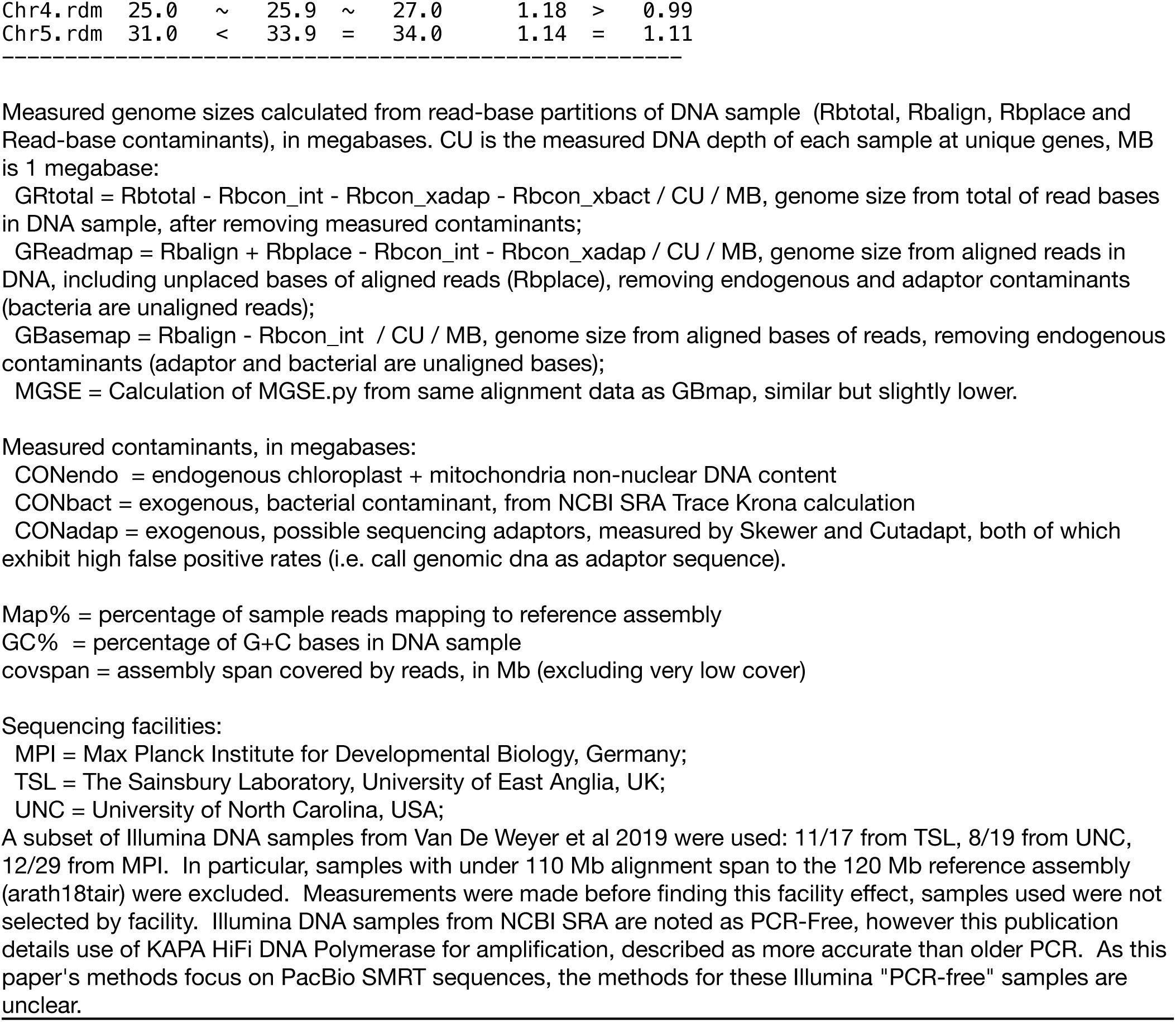
Sequencing facility methods effects (A) on measures of DNA samples of *Arabidopsis* ecotypes (Van De Weyer et al 2019), and chromosome assembly (B) effects. Values are megabases averaged over ecotype samples, or percentage for Map, GC, +/- S.E.M. The MPI samples are significantly different from the other two (TSL, UNC) for all measures: reduced genome sizes, increased contaminants, lower map rate, and higher GC content. Assemblies are those of Gnodes#1 Table R6.1, measured with TSL DNA samples. See supplemental tables gnodes2tab4_at19eco36info.txt, gnodes2tab4_at19eco36gnodes.txt for further details.

Table T4 also indicates that laboratory methods can have a large effect on genome measurements. In this example, one of three sequencing facilities produced samples with significantly less genomic DNA, and with more contaminants. The other two facilities produced DNA samples that are nearly the same, on average, and that match flow cytometry measures for this species (166 mean, 160-172 range by FC of Long et al 2013 versus 164 mean, 156-172 range of Gnodes GRtotal). The biosamples differ among facilities, but each facility’s samples span the species range in Europe.

Table T4 (C) has measured DNA sizes of chromosomes for three assemblies across ecotype samples. Whereas DNA size of full genome is nearly same across assemblies, chromosomes differ somewhat, due to where sample DNA is mapped. The 11 ecotype samples from TSL facility have a small range with low variance. Note at18tair_Chr3 has 10 Mb higher est. size, 15 Mb above observed size, but lower est. for other 4 chrs, largely an effect of centromere DNA of all 5 chrs (2 Mb each) mapping to one Chr3 locus, non-existent in other chrs of this assembly. Centromeric DNA is mapped evenly across chrs in other two assemblies with centromeres. Estimated chrs sizes are statistically same for at21ncbi and at22canu, but for Chr4 which may be due to extra rDNA spans of at22canu assembly (see Figure R6.1). DNA of another ecotype was sequenced with PacBio HiFi and Illumina PCR-Free methods, and its measured readmap sizes are within statistical range of samples, but for Chr5 which appears lower. PacBio HiFi and Illumina PCR-Free results are same within stat. error, differences may be due to mapping effects for different read types.

#### S2h. Computational errors in error-correction, filtering

Long-read error correction methods, as used in DNA assemblers, reduce duplicated and repeat regions to averaged or filtered subsets. Three corn isolines (Hp301, NC350, Ki3), with Pblo DNA (data of Hufford et al. 2021) were processed with Canu error correction step. These three had Cym and DNA genome size estimates in the 3,000 Mb range, well above the 2,200 Mb assembled reference. After error-correction, the DNA estimates were reduced by one-third or more (hp301 3200 > 2200, nc350 3150 > 2250, ki3 3600 > 1700), reductions in duplicated DNA, TE and repeat content.

Another long-read error correction method, of the Marvel assembler (Nowoshilow et al 2018) developed for the highly repetitive axolotl genome, was tested on Pblofi DNA of the fire ant, which has large amounts of high order repeats (see Gnodes#3). Canu error correction reduced input DNA by 54%, while Marvel reduced it by only 14%. Both of these reduced DNA size measures from the original, Marvel was less drastic: 15% reduction in total, mostly in the duplicated repeats and TE contents, while Canu correction reduced size estimates by 28%, below fire ant assembly size.

Averaging and filtering of repetitive DNA is important for genome assembly; it also affects completeness, and is a difficult problem for genome informatics. When natural copy numbers of repeats are reduced artificially, that can impact assembly of these regions, leading to the observed Cym - Asm discrepancy.

Long-read high fidelity data types (PacBio hifi or CCS, ONT duplex) achieve a high accuracy relative to assemblies of those data, see Figure F7.MErr. This is done with computational methods that post-process the machine outputs, beyond base-calling. Both types rely on software that uses methods of averaging or filtering the machine read outputs. Current PacBio software for producing CCS (Hifi) samples has computational options that change how it will recover all the DNA of the samples, whereas earlier CCS software and default parameters appear to bias DNA hifi sample toward non-repetitive fractions in the quest for higher accuracy.

The sample of two ONTduplex data sets (Koren S et al. 2024) is too small to draw firm conclusions, and software to accurately process ONTduplex is in flux. One conclusion is apparent: ONTduplex quality results for human and corn are quite different. The human result matches current genome assemblies from other sources. The corn result, while matching large parts of others, is deficient in spans of rRNA, satellite repeats and some others. This might be explained by a low duplex processing rate obtained for corn of 16%, versus 55% for human (Koren S et al. 2024, Suppl Table 1 Human HG002 cell line Duplex sequencing and Suppl. Table 5. Maize Duplex sequencing). This poor result for corn ONTdp may be a correctable artifact of laboratory or DNA processing methods.

Gnodes #1 (Gilbert 2022) documented under-estimates of genome size by GenomeScope and other k-mer methods, due in part to use of truncated k-mer distributions, a result of poor defaults in genome software and/or assumptions that very high copy duplications are contaminants or artifacts. Use of truncating default options for DNA k-mer measuring software cuts their estimates substantially (Arabidopsis: trunc 125 vs full 151 Mb, Daphnia: trunc 137 vs full 242 Mb; Gnodes #3, Table 3C, Gilbert 2023). Care must be taken to collect a complete k-mer distribution; a recent Arabidopsis publication (Lian et al. 2024), with a "high" cut-off of 2 million still truncates the full distribution by 3%-5% of genome sequence.

MarkDuplicates is a common tool for PCR duplicate removal, but it does not differentiate natural and artifact duplicates, relying on reads mapped to an accurate chromosome assembly. As assemblies are often inaccurate in duplicated regions, such duplicate removal is problematic: measured on PCR-Free DNA samples of Illumina and Nanopore, MarkDuplicates marked as (artifactual) duplicates large portions (12% - 20%) of arabidopsis and corn DNA from natural parts of these genomes. This agrees with findings of Bansal (2017) who reports that 50% - 90% of reads marked as duplicates are natural, not artifact. For DNA samples with PCR amplification, this work recommends no duplicate removal for measuring genomes, as a large portion will be natural, and the artifactual PCR additions, if quasi-random, will not as greatly affect measurements. When particular components of a genome are over- or under- amplified, biases are a problem (Gnodes#1 Table R8.1, Figure R8.1); the best choice is to avoid using DNA samples produced with possibly biased amplification methods.

There is a similar artifact versus natural trade-off with sequencer adaptor trimming software, including CutAdapt and Skewer, that use error-prone alignment to known adaptor sequences, with relatively high rates of false positive matches (e.g. the most common, 3-base matches are unsupportable), and call adaptors at rates in reference genome assemblies similar to high quality DNA read samples. For measurement use, it is recommended to not preprocess DNA samples by trimming adaptor-like sequences, but to separately measure such as desired. In good samples, 0.1% to 0.4% of DNA is called as such (Table T4).

The computations used in genome assembly are not a focus here, however the problem of distinguishing biological from artifact in high identity duplications are fundamental to precision genome data.

Accurate measurement of DNA contents is an important step in determining quality of genome assembly. With such measures, the value of assembly tools can be identified. Natural variation among duplicate sequences of different loci has similar signals to machine error and same-locus heterozygosity. For example, Jaegle et al. (2023) find that most "heterozygous" classed SNPs in *A. thaliana* are artifacts of duplicated regions, rather than residual heterozygosity. Among assembly methods with problematic filtering results are the "heterozygosity" filters, such as purge_haplotigs and purge_dups. As reported in Gnodes#3 (Table 3B), these programs remove significant fractions of human and plant reference genomes, in regions of high satellite DNA content, purging 60% of human Y, 10% of At_plant Chr2.

Table T6 read_mappers_long_short has summaries from Gnodes for SR and LR read sets of five well studied organisms (model plant, beet, chicken, human and corn). Corn and beet plants are especially informative as they have high rates of duplicated genome, notably more than other organisms measured. Summary statistics of genome size estimate from total read bases and mapped bases, portion of bases aligned, unplaced and unmapped, genes classed by DNA depth as unique, duplicated or poorly measured, and genome coverage and depth found for the read set/mapper.

minimap2 and bwa-mem with short-read data produce similar results, a small improvement for bwa in read-map rate is balanced by slightly higher map error rate; likely options for these two can be adjusted to increase SR map rate or reduce error rate. minimap2 for long read data performed well, near equivalent to short-read for the more accurate HiFi LR data. winnowmap2 is closely similar to minimap2 (which it uses for alignments, following k-mer analyses), and has small improvements for some read sets over minimap2, but winnowmap2 commonly underperforms in alignments to gene sequences, with a high rate of zero or skew gene coverage. Long-to-short split reads had mixed results, with minimap, the more accurate HiFi when split were measured to be near equivalent to short read alignments, but the less accurate Nanopore and lofi PacBio had poor results with split to small pieces, notable with rather low map rates. lra performed poorly for Gnodes use, with notably greater unmapped reads and incomplete coverage, on both chromosomes and gene sequences, and larger variance in measures. minimap2 is chosen as default aligner of both SR and LR data for Gnodes use bases on these results.

#### S2i. Do DNA and cytometric measures agree on genome sizes?

The best evidence for agreement of DNA read measures with other genome size measures are studies using the same bio-samples, controlling biology, laboratory effects as much as feasible. A few such studies have been undertaken, notably with At model (Long et al 2013) and corn plants (Hufford et al., 2021). These are limited and ambiguous; the early At Illu DNA sequences were very biased in GC content, likely due to PCR chemistry. Corn plant results (Table T2B) show a close agreement between Cym and long-read DNA measures of size, in average size (3% difference) and a 98% correlation over 8 isolines.

A lab effect on GC content of DNA was noted (Table T4A). The GC content of current full assemblies of At plant is 36-37%, but DNA of the MPI facility has a 4% higher bias, about the same as for genome size measure, 3.1%. An extreme case of GC bias in DNA reads is the valuable but unreplicated work (Long et al 2013) on At ecotype lines that measured both Cym and DNA contents for 100s of lines in a geographic range, finding a cline in genome sizes consistent between Cym and DNA. The GC bias of this Illumina DNA is 6%, which obscures measures of ecotype genome variation, as this amplification bias differentially affects CDS, transposons, rRNA and other duplications. This work, unfortunately, has been used in benchmarking and validating DNA measurement software.

The discrepancy of techniques for genome measurements has been attributed to cytometric methods producing an over-estimate bias (eg. Sun et al 2017, findGSE), but the argument presented for this is weak: no evidence for an over-estimate bias by Cym is produced, instead the authors argue that since k-mer DNA methods match assembly sizes, they must be correct. Under-estimates by Cym are probable, if the main error factor is prior size estimates of standards (eg. Temsch et al. 2022). Indeed this corn isoline report (Hufford et al., 2021) uses Cym under-estimated by 10-15%, compared to the Cym data reported in the cited references (Chia et al 2012, cited by Wang et al 2021 and Hufford et al 2021) [details : Chia et al 2012, Suppl. 1.13 flow cytometry: used chicken standard, approx. 50% of corn genome size, and measured corn CE-777 line as 2580 Mb (5.28pg/2C), consistent with other corn measures; Wang et al 2021 use these values, and also measured corn B73 line as 2550 mb (5.23pg/2C) by Cym. However they then use 2300 mb for the Cym size in analyses, and Hufford reports 2100 mb as B73 line estimated size, i.e. a 10%-15% underestimate of their Cym values]. Pervasive under-estimates of genome size by common DNA processing, measures and assemblies are detailed here, as well as elsewhere (Mgwatyu et al. 2020).

To resolve this problem of accurate genome size measurement, an effort to reproduce cytometric and DNA measures with careful laboratory techniques, using species samples that control as many biological effects as feasible, is warranted. The approach used by prior studies of inbred lines known to vary in genome size, for At and corn plants, seems suited to this. From the results presented here, evidence supports that, for the least-biased DNA samples of Oxford Nanopore technology, they agree with cytometric measured genome sizes.

## Discussion

"If you can’t measure it, it isn’t real" (Lord Kelvin, scientist in 19th century) applies to genomes: if you can’t measure it accurately, it isn’t really true. We would like to know how real are genome assemblies, more accurately called pseudomolecules in some literature.

The noted discrepancy of genome sizes, between cytometry and DNA assembly, can be attributed to various DNA processing steps that reduce duplicated genome content. Results reported here are conclusive that several DNA processing biases reduce duplications. DNA per-base error correction methods are collapsing natural duplication variance into averaged sequences. DNA filters applied for presumed complications such as heterozygosity and artifacts are reducing or ignoring natural duplicated DNA. The results presented here show that the least filtered, or corrected, long read DNA will measure genome sizes that are closest to that of other methodology (Cym), and have more agreement among measures. This doesn’t exclude errors in cytometric methods, but no evidence of a systematic measurement bias for those has been established (Temsch et al. 2022).

A review of sequencing and methodological problems with duplicated and repetitive DNA (Cechova 2021) has identified several sequencing technology factors that affect these measures, indicating that each method has its own problems and values, but in general they agree with each other, and are all "probably correct". As indicated in this report, agreement at or below 90% is established, the difficulty for precision is assessing an uncertain 10% in measures.

Recent Arabidopsis ecotype data (Lian et al. 2024) are consistent with the overall results reported here, in Table T2prime, an update of Table T2. Ontl is closest to cytometric genome size, with lowest variance, while Pbhi tends low, and Pblo tends high. Short reads have a larger range of results.

**Table T2prime.** Genome sizes (Mb) from short & long DNA, averaged over Arabidopsis ecotype lines [Bioproject PRJEB62038 of 2024], and long - short differences. Long read types (Ontl24 or Pbhi24) were produced at different labs, from different but related ecotypes. Short read samples are paired with long types for each sample. Pblo19, Illu19 are of Table T2. Coefficient of variance, CV, is given.

| Item | Pblo19 | Ontl24 | Pbhi24 | D(0-P)24 | Ont24 | Pbh24 |
| --- | --- | --- | --- | --- | --- | --- |
| LongRead | 162 | 155 | 147 | 7.9* | CV 2.6 | 4.0 |
| ShortRead | 156 | 149 | 157 | -8.7* | CV 6.0 | 3.4 |
| D.lr-sr | 6 | 6* | -10* | *p < 0.005 |  |  |

Significant differences exist between long - short of each lab group, and between long types from different labs or ecotypes. These differences are likely due to differing laboratory methods (see Table T4A), as well as sequencing technology.

### Recommendations for measuring genome sizes and contents

Use Ontl, as least biased long-read type, to measure genome sizes and contents most accurately. Use two DNA types to estimate the range in measures.

Use cytometric measures, quantitative PCR and/or other types of genome size measures for comparisons.

Measure contents during steps in DNA filtering for assembly. Comparison of unfiltered versus filtered DNA, with assembly contents, is a quality check for completeness of genome assembly.

## Conclusions

Genome sizes from assemblies of animals and plants are often smaller than cytometric measures. A third measure, from DNA sequence samples before assembly, divided by the unit depth of conserved unique genes, generally support the larger cytometric measures.

## Methods

The major contents of genomes presented in these results are computed with Gnodes, as described in Gnodes#1 (Gilbert 2022). Gnodes software is part of the EvidentialGene package, publicly available at http://eugenes.org/EvidentialGene/ and http://sourceforge.net/projects/evidentialgene/ [Gilbert, 2019].

The basic algorithm of Gnodes is (a) align DNA reads to gene and chromosome assembly sequences, recording all multiple and unique map locations, (b) tabulate DNA cover depth at each sequence bin location for multiple and unique mappings, (c) measure statistical moments of coverage depth per item, itemized by categories of gene-cds, transposons and repeat annotations, duplicate and unique mappings, (d) summarize coverage and annotation tables, at chromosome-assembly and gene levels. The statistical population of coverage is non-normal, so that median, average, skew and other statistics are calculated to approach precise and reliable measures.

### M1. Data sets: DNA read sets, genome assemblies & annotations

DNA reads are drawn from public archives at NCBI SRA, and most chromosome assemblies from NCBI Genomes public archive. Supplemental tables for this paper include **gnodes2dnatable.txt**: SRA IDs of genomic DNA used for each species assembly, with count, read technology type and average read length; **gnodes2chrasm_idtable.txt**: genome assembly IDs, size, date used per species; **gnodes2chrasm_metadata.txt**: genome assembly meta data as used for Gnodes, including ID, Species, size, GenbankID or assembly URL, Date.

Gnodes summary analyses of these DNA are tabled in **gnodes2allspecies_sum.txt**, which includes columns

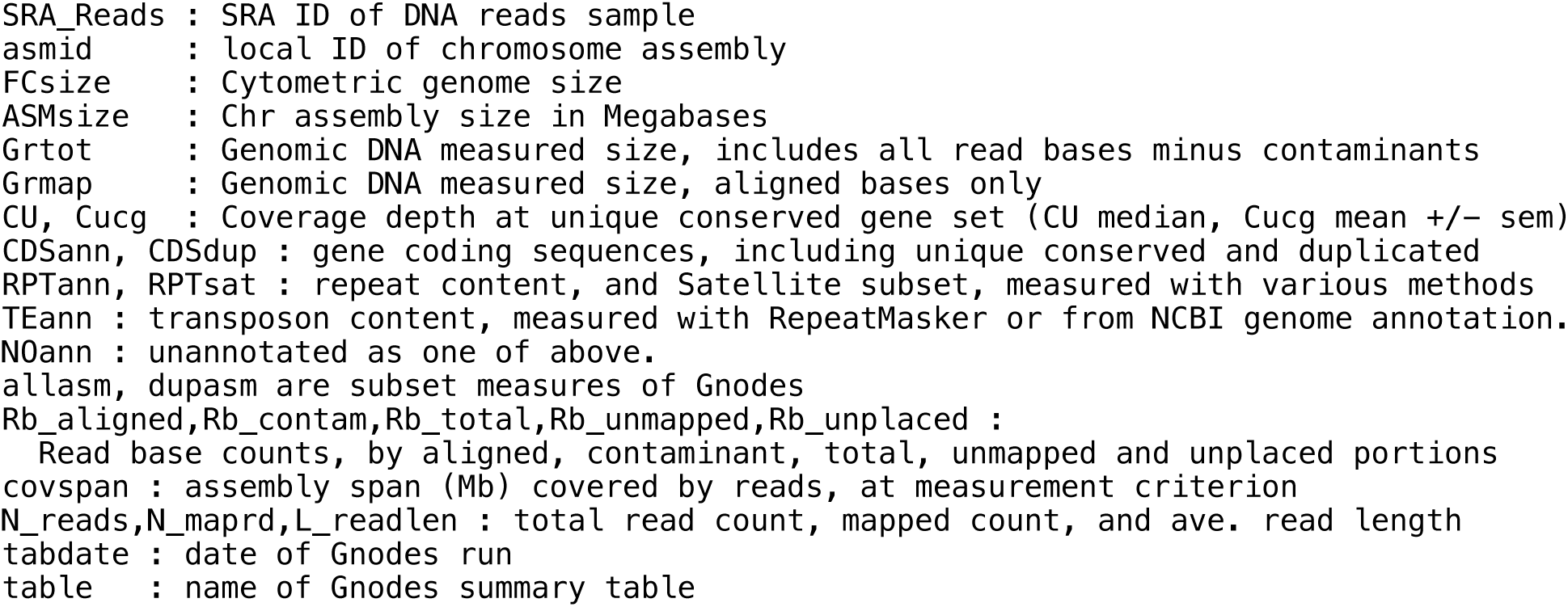

**gnodes2allnonmim_sum.txt** has Gnodes summary data for non-mimimap mapping methods, where SRA_Reads ID and asmid match those of gnodes2allspecies_sum.txt done with minimap2.

#### M2. DNA mapping methods

The author has examined several long-read DNA mapping programs: minimap2 [doi: 10.1093/ bioinformatics/bty191], widely used, is the default mapping software for Gnodes. For short-read mapping, bwa-mem2 [arXiv:1303.3997] and minimap2 are used. Others tested that build on minimap2 include (1) Winnowmap2 (Jain et al. 2022 doi:10.1101/2020.11.01.363887v1), which adds k-mer analysis of repetitive DNA in mapping. Comparisons done for this project found winnowmap produces small differences from minimap2 which were inconsistent in terms of accurate measurement of repetitive content. (2) blend (Firtina et al. 2022), also based on minimap2, which its authors recommend for pacbio-hifi DNA. Tests for this project found that blend had lower hifi alignment compared to minimap, with more unplaced and unmapped reads. (3) lra (Ren et al. 2021 doi: 10.1371/ journal.pcbi.1009078), a long read aligner, which did map long reads, but poorly for this author; (4) TandemTools (Mikheenko et al. 2020 doi: 10.1093/bioinformatics/btaa440), which failed to function for this author. As a test, long reads were mapped as short ones, by cutting to paired short-read segments, with poor results.

#### M3. Genome annotations

Sequences of reference annotations for species were used by preference, where available in good quality, for coding genes, transposons, and repeats. Reference gene CDS were extracted from gene data provided by NCBI Genomes [ncbi-genomes/GCF_accession*rna.gbff]. The outputs of RepeatMasker, and RepeatModeler for some cases, were used [Smit et al, http://www.repeatmasker.org]. Computations done for this project include runs of tandem repeat finder (TRF, Benson 1999) and high order repeat finder (SRF, Zhang et al 2023) for some genome assemblies. Annotations used with the Maize NAM isoline assemblies were produced for each isoline, in preference to use of reference Maize assembly, in accordance with the large isoline differences in genome contents.

#### M4. Major genome content measures

Genome assemblies and major contents sequences of CDS, transposons, and repeats were processed with Gnodes, which uses BLAST alignment of these to produce genome location tables congruent with the genome read-coverage tables also produced by Gnodes. Component scripts of Gnodes summarizes and plots DNA copy numbers relative to genome assembly (xCopy), and megabase measurements for the major contents.

#### M5. Methods for figures and tables

Figure F1.sum. (A) Dna and Cym sizes are from gnodes2allspecies_sum.txt, subsets by read type. (B) Genome sizes for animal and plant species, as Assembly / Cytometric size percentage in relation to cytometric sizes. NCBI Genomes (NCBI 2023) provides assembly sizes, median value for multiple assemblies, from year 2020 to 2023, many of long-read DNA methods. Animal Genome Size Database (Gregory 2023) and Plant DNA C-values Database (Leitch et al 2019) provide the most recent cytometric sizes of the same species. See Gnodes#3 (Gilbert 2023) for supplemental details.

Figure F2.rRNA. Human and corn rRNA gene copy numbers over gene spans, from Gnodes.covtab / CU and rRNA gene annotation locations.

Figure F3.agr and F6.agrd : agreement statistics by read type and for major contents of genomes. Methods described in S2e. Agreement for high copy genome contents.

Tables T1, T3. Re-map rates and contents of long-read unplaced spans: Unplaced sequence spans, 150 bp or longer, from long reads that partly map to chromosomes were extracted and re-mapped to assemblies, using both blastn and minimap2. Bases that re-align to assembly are tabulated, including their annotated assembly content types.

Figure F1.sum.D : Major contents as percent of assembly, by read types, as stacked bar plot. Figure F2.rRNA.C,D, F4.TEcn : extension of F2.rRNA.a,b.

Figure F5.TEx.A-F : copy number over TE example spans, as in F2.rRNA.a,b, for transposons. Figure F7.MErr : map errors for read types, calculated by Gnodes from read x assembly mapping tables.

Table T4. Sequencing facility methods effects [Table T4 has methods footnote.] Table T5. Chromosome parts measures of At. assemblies.

Table T6. read_mappers_long_short [need some methods explained]

**Table T5.**
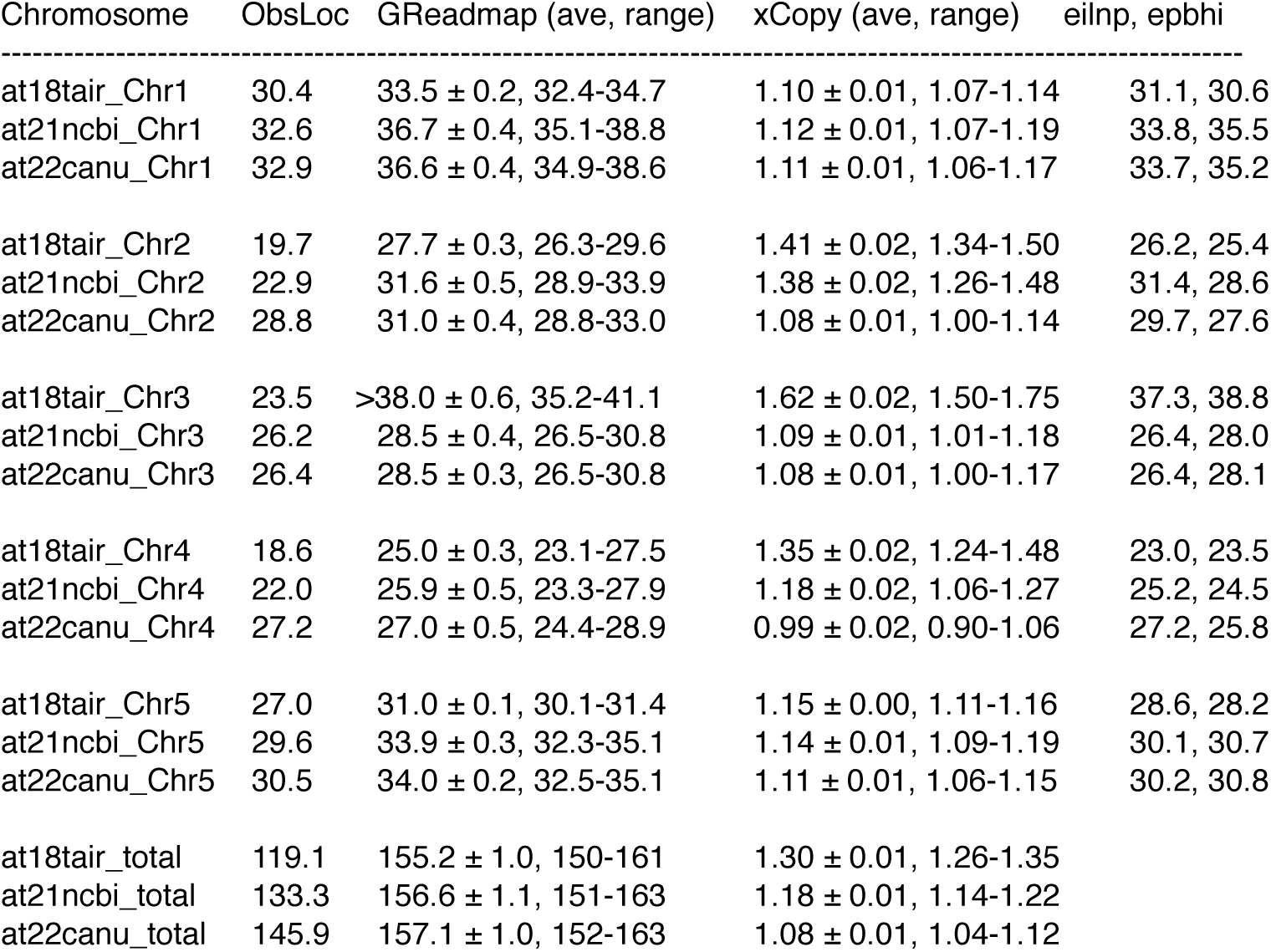
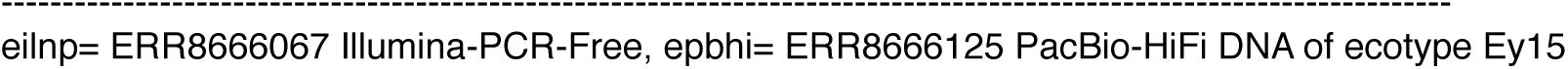
Chromosome parts measures of At. assemblies, for 11 DNA samples of Arabidopsis ecotypes of TSL facility, and for Illumina-PCR-Free (eilnp), PacBio-HiFi (epbhi) of ref ERR8666, recapping Table R6.2, 6.3 of one DNA sample. ObsLoc is the observed assembled size. GReadmap is estimated size from DNA read samples, +/- SEM and sample range, xCopy is est./obs. size ratio.

**Table T6.**
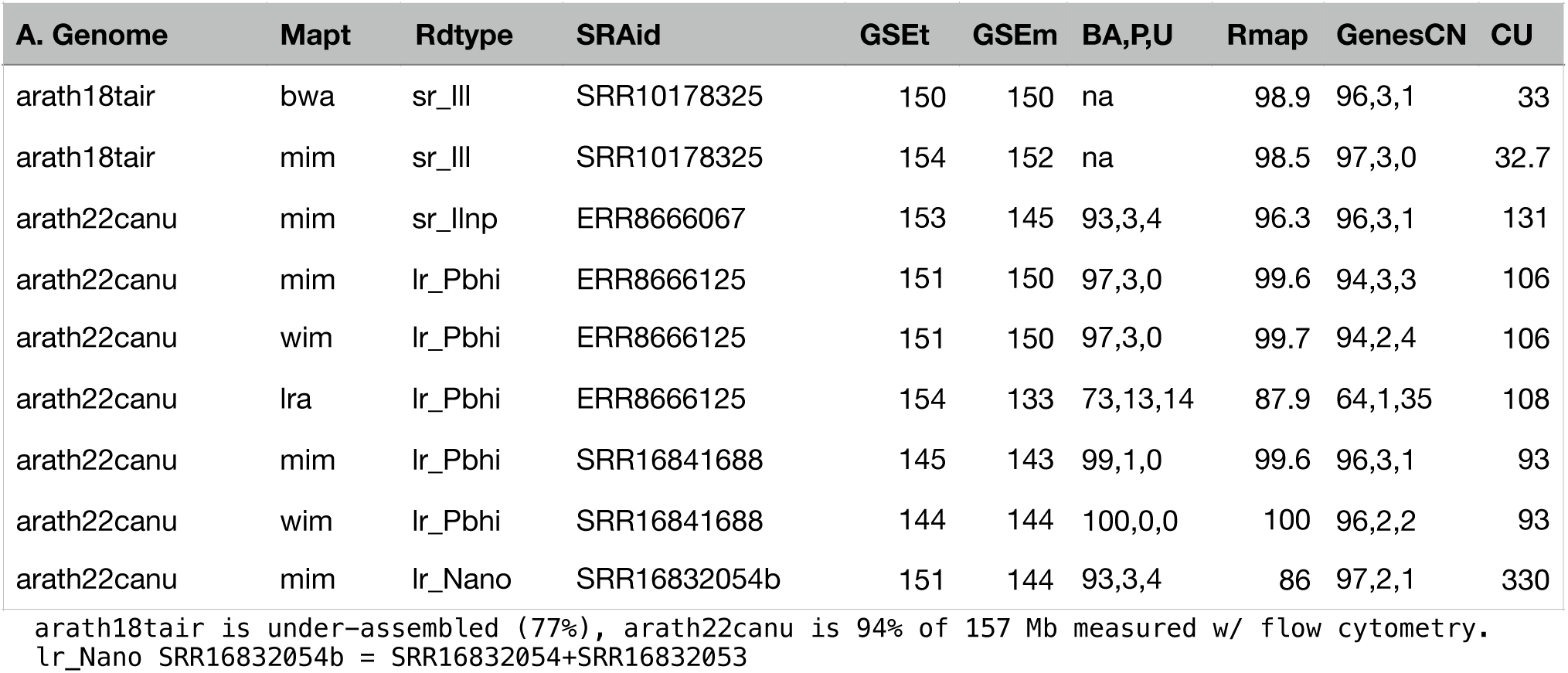
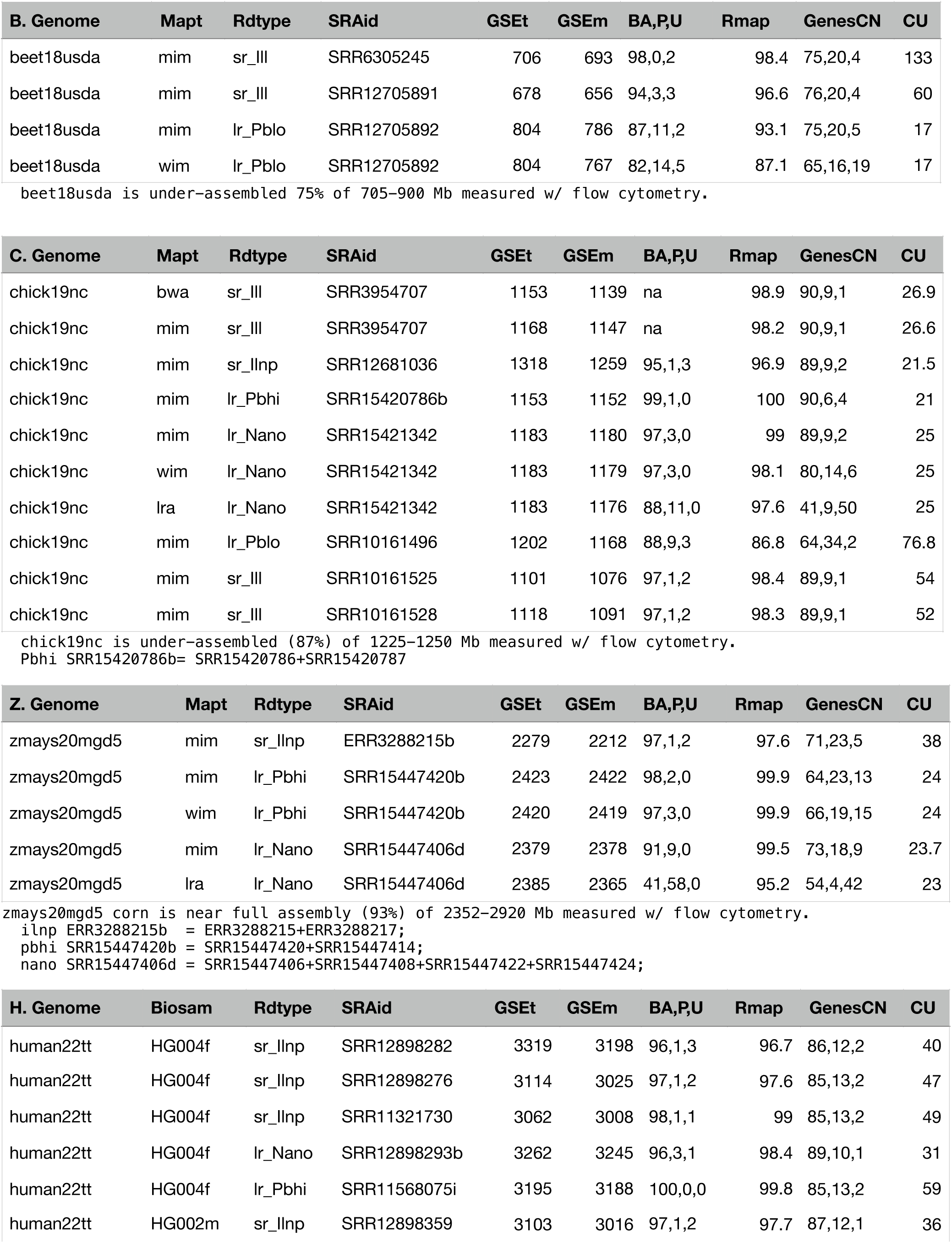
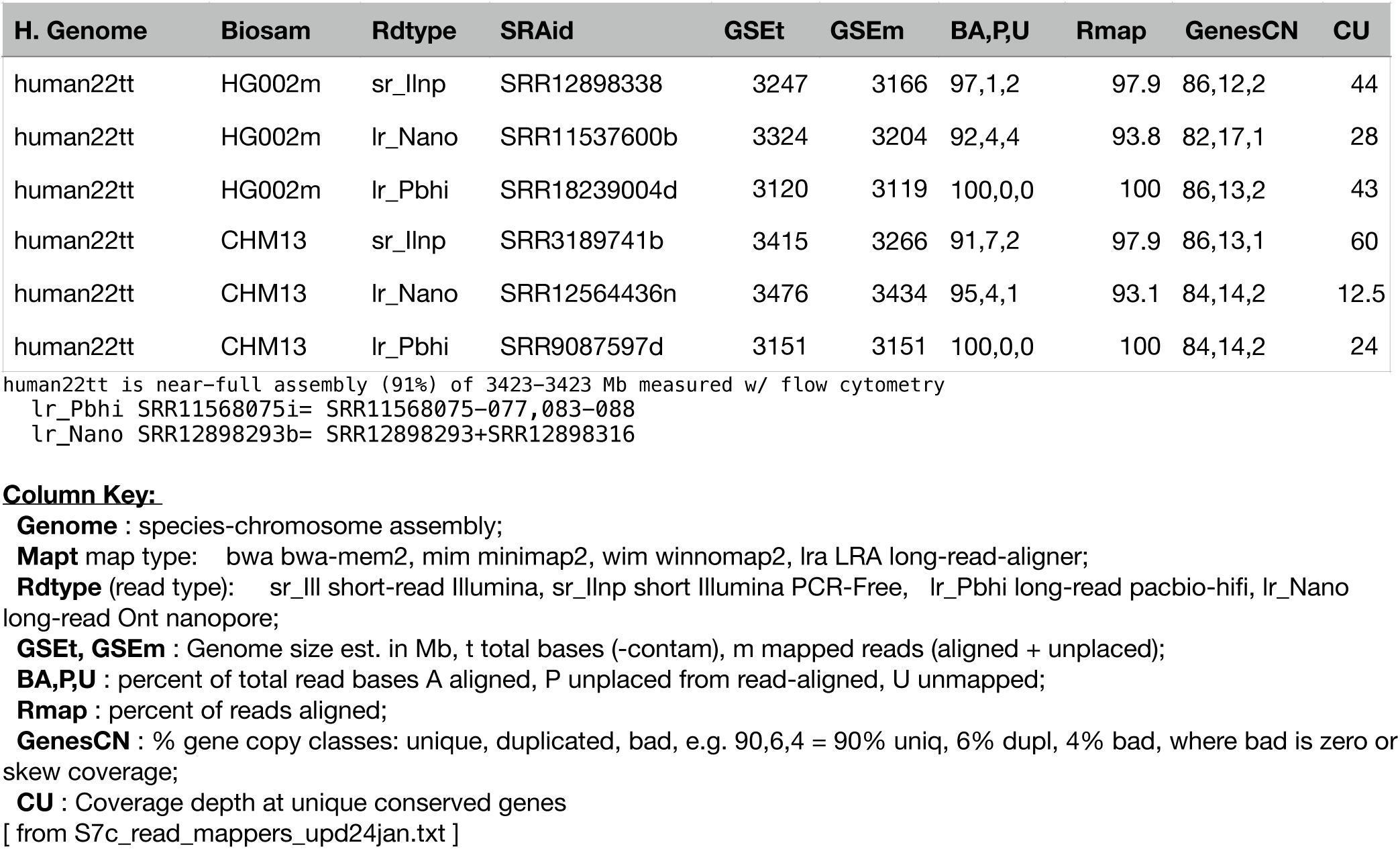
read_mappers_long_short. Read map software, with long and short read samples aligned to chromosome and gene sequences, are compared for use in Gnodes quality measures and genome size estimations. Genomes measured are A. *Arabidopsis thaliana*, B. *Beta vulgaris* (sugar beet), C. Chicken (*Gallus gallus*), Z. *Zea mays* (corn), and H. *Homo sapiens* (human). Read types are Illumina paired short read (SR), with PCR-Free methods as available, and long reads (LR) of PacBio HiFi and lofi, and Oxford Nanopore.

## Acknowledgements

The following provide shared computational resources in support of this work: Indiana University Research Computing, Extreme Science and Engineering Discovery Environment (XSEDE) then Advanced Cyberinfrastructure Coordination Ecosystem: Services & Support (ACCESS), supported by National Science Foundation, including Jetstream (jetstream-cloud.org) and San Diego Supercomputer Center.

